# Novel PFAS in Alligator Blood Discovered with Non-Targeted Ion Mobility Spectrometry-Mass Spectrometry

**DOI:** 10.1101/2025.03.20.644452

**Authors:** Anna K. Boatman, Gregory P. Kudzin, Kylie D. Rock, Matthew P. Guillette, Frank Robb, Scott M. Belcher, Erin S. Baker

## Abstract

Per- and polyfluoroalkyl substances (PFAS) are a large and growing class of chemicals gaining global attention due to their persistence, mobility, and toxicity. Given the diverse chemical properties of PFAS and their varying distributions in water and tissue, monitoring of different matrices is critical to determine their presence and accumulation. Here, we used a platform combining liquid chromatography, ion mobility spectrometry, and high-resolution mass spectrometry for non-targeted analysis to detect and identify PFAS in alligator plasma from North Carolina (5 years, 2018-2022) and Florida (2021 only). Structures for 12 PFAS were elucidated, including 2 novel structures, and an additional 34 known PFAS were detected. Three of these compounds were previously unreported in environmental media. More PFAS were detected in North Carolina alligators and no novel PFAS were detected in Florida gators. Quantitative analysis of 21 of the known PFAS revealed that plasma concentrations did not change over the 5-year study.

**Highlights:**

- 46 PFAS were detected in blood plasma from North Carolina and Florida alligators using LC-IMS-HRMS
- Structures for 12 previously unreported PFAS were elucidated, including 2 novel structures
- Cape Fear River alligators had the most types and highest concentrations of PFAS

## Introduction

Clean drinking water free of unwanted and harmful chemicals is a basic human right. Per- and polyfluoroalkyl substances (PFAS) have gained widespread attention because of their unparalleled persistence, mobility, toxicity, and presence in humans, wildlife, and ecosystems globally around the world. Some individual PFAS have been well studied and regulation is increasing; however, PFAS are a large and growing class of chemicals, and the presence and health impacts of most of them are not yet understood. Detecting unknown PFAS is therefore a critical part of determining their impact on human and ecosystem health. To date, an incredible amount of research efforts have focused on drinking water and surface water using both targeted and non-targeted analyses.(*1-5*) Targeted analyses with mass spectrometry use analytical reference standards to quantify known PFAS in water and biological matrices, but are inherently limited to only a few dozen PFAS. When standards are not available or investigations into new PFAS are desired, non-targeted analyses (NTA) are used. NTA efforts in water and biological tissue have revealed that different PFAS are detected in different matrices(*6-8*), underscoring the importance of testing both the exposure source and the exposed organism.(*6, 9*) However, NTA of complex biological matrices is challenging. Obtaining biological samples often requires time- and resource-intensive collection processes. Moreover, these samples have many other molecules present, such as proteins, lipids, and metabolites, resulting in feature lists dominated by highly abundant biological molecules with no relevance to the analytes of interest. Work to improve and increase NTA analyses in biological samples is of the utmost importance for contaminant studies, including PFAS.

The lower Cape Fear River (CFR) in North Carolina serves as a primary drinking water source for local human and animal residents and is famously located downstream of a fluorochemical manufacturer that produces Nafion using perfluoroether compounds. Previous non-targeted studies in this region discovered perfluoroethers in seabirds(*6*) in addition to other unknown fluorochemicals in water(*5*). Targeted, quantitative studies in the CFR have also consistently noted high PFAS levels in water(*1, 5, 10*), fish and other wildlife(*6, 11*), human serum(*4, 12*), and domestic sentinel species including horses and dogs.(*13*) In their targeted study of alligators sampled in 2018-2019, Guillette *et al*. found associations between increased serum PFAS levels and several immune health endpoints, including increased autoantibody presence and field observations of unhealed wounds. Reanalysis of the same alligator serum samples by Bangma *et al*. showed the presence of additional perfluoroethers for which standards had not been available in the original targeted analysis.(*9*) Alligators are valuable sentinels for human exposure to harmful chemicals due to their long life span and position at the top of the food chain, both of which result in high levels of exposure to aquatic bioaccumulative pollutants. Additionally, their complex and robust immune systems make them sensitive predictors for exposure to immunotoxic chemical pollutants such as PFAS. The study authors concluded that the presence of these compounds in alligators is cause for concern about human exposure, and that the detection of additional PFAS with no standards available means that reported PFAS levels underestimate the total amount of PFAS present. Their results clearly demonstrate a need for ongoing biological monitoring of PFAS levels in people and animals from this region.

In this study, we contribute 5 years of alligator PFAS NTA data (2018 to 2022) to the growing body of PFAS research on CFR water and wildlife. To contrast with the CFR, we included alligators from Lake Waccamaw (LW), a nearby reference site located within the Lumber River basin in North Carolina, as well as alligators from St. John’s River, Florida (FL), a geographically farther away site with no known PFAS industry nearby.(*14*) The primary aim of this study was to identify as many PFAS as possible in alligator plasma with a focus on novel and uncommon compounds that could go undetected in NTA of water. We additionally quantified up to 26 known PFAS to evaluate geographic and temporal exposure trends. The combination of quantitative data and information on novel PFAS will be of great interest to researchers and local residents because these results uncover previously unreported PFAS that have accumulated to detectable levels in a sensitive sentinel species.

## Materials and Methods

### Study Area and Sample Collection

All animal procedures were performed with approval by the North Carolina State University Institutional Animal Care and Use Committee (protocol #18-085-O). Alligators were sampled using active capture methods described previously(*15*) and animals were released at the site of capture. Sampling locations included several sites within the lower Cape Fear River basin in North Carolina, one nearby reference site, Lake Waccamaw, in the Lumber River basin, and one site in Florida (St. John’s River) (**Figure 1, Data S1**). Sampling occurred between July 24, 2018 and August 17, 2022.

Blood samples were collected from each animal immediately following capture as described previously.(*15*) Plasma was collected in lithium heparin coated plasma tubes (Vacutainer, BD, Franklin Lakes, NJ), stored on ice until centrifugation (1800x g for 10 min at 4C), and then stored in polypropylene microcentrifuge tubes at -80C until analysis. Serum was used in the original alligator study from 2018-2019; however, plasma was used in this study due to its easier handling and higher yield volumes in the field compared to serum. Preliminary proof of concept work (not published) comparing alligator serum and plasma indicated no significant differences in PFAS detection between the two matrices.

**Figure 1.**
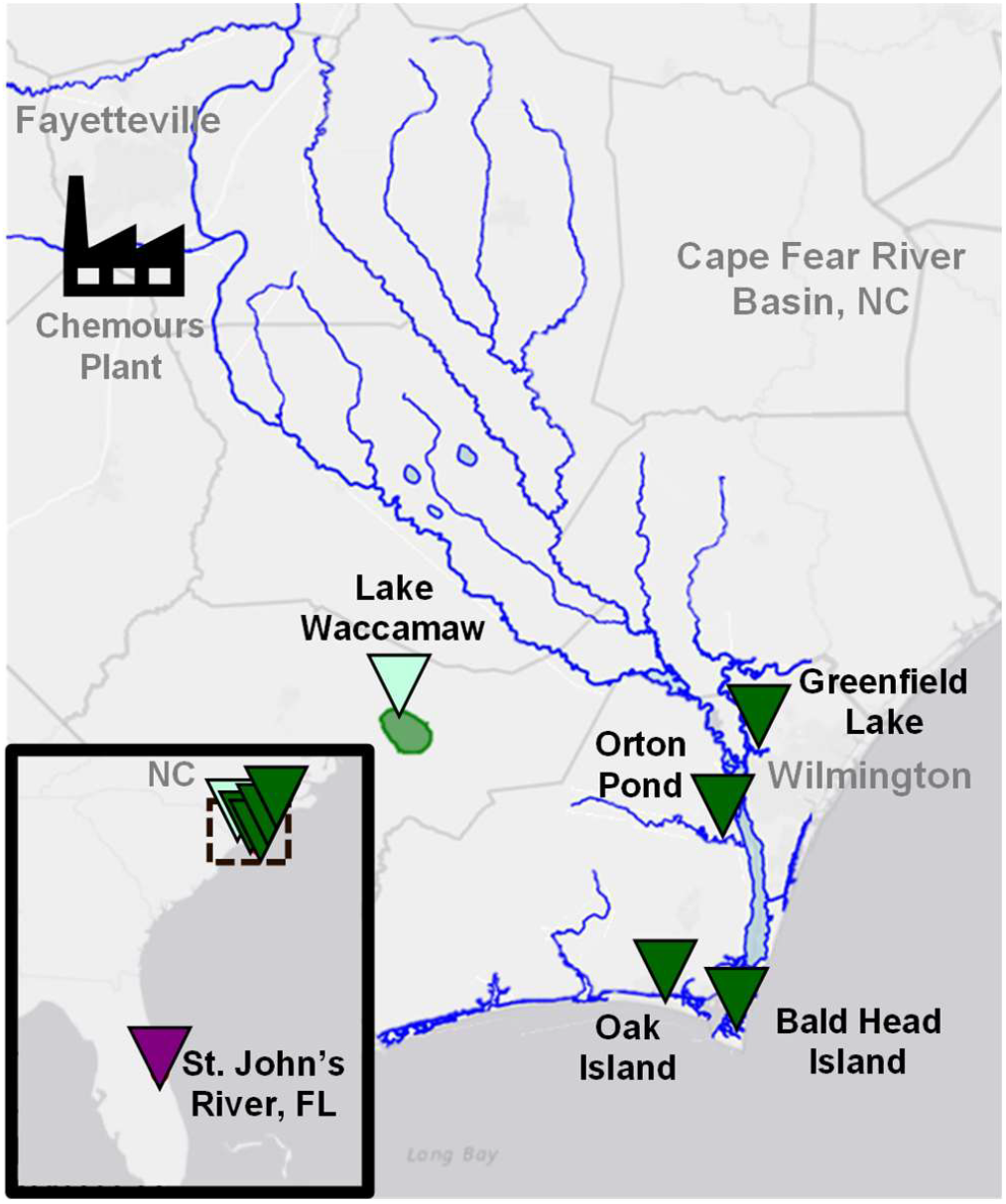
Map of alligator sampling sites. Sites are shown in relation to the Cape Fear River basin, NC and the upstream Chemours manufacturing facility. CFR sites are indicated with dark green triangles, LW with light green, and FL with dark purple.

### Standards and Reagents

A heavy-labeled PFAS standard mix (MPFAC-HIF-ES, Wellington Laboratories, Guelph, Canada) containing 26 different 13C- or D-PFAS at concentrations from 250 to 5,000 ng/ml was diluted 1:1 in methanol for use as internal standards. The Standard Reference Material 1957 (SRM1957, human serum) was obtained from NIST (Gaithersburg, MD) and used for quality control. Calibration curves were prepared in charcoal-stripped fetal bovine serum using seven concentrations of native PFAS standards (PFAC-MXH and HFPO-DA, Wellington Laboratories) ranging from 0.128 to 4,000 ng/ml and a constant concentration of the heavy-labeled internal standard mix. All solvents (acetonitrile, water, and methanol) were Optima LC-MS grade from Fisher Scientific.

### Plasma Sample Preparation

Alligator plasma samples were blocked and blinded into four batches for extraction. A 50 µl aliquot of each sample was transferred into a polypropylene microcentrifuge tube, along with three field blanks and three QCs (NIST SRM 1957 human serum) per extraction batch. Full extraction method steps are detailed in the **Supplementary Materials**. In brief, internal standards (5 µl) were spiked into each aliquot, proteins were precipitated using cold acetonitrile (300 µl), and a portion (200 µl) of the acetonitrile layer was evaporated to dryness. Samples were reconstituted in 40:60 methanol:water buffered with 3 mM ammonium acetate (100 µl) and transferred to polypropylene LC vial inserts. Pooled alligator plasma samples and matrix-matched calibration curve samples, along with three method blanks and three QC samples, were prepared following the same procedure in a separate batch.

### LC-IMS-CID-HRMS Data Collection

All samples were first analyzed on the LC-IMS-CID-HRMS platform in MS1-only mode (no collision energy).(*16*) Samples were blocked by sample preparation group, site, and year and then randomized to mitigate the potential for introduced bias from the worklist order. The pooled samples and several representative individual alligator samples were reinjected using the size-dependent AIF method described previously by Donelson *et al*.(*3*) Analytical methods are summarized briefly here and full details are provided in the **Supplementary Materials**. A C18 column was used (Agilent Zorbax Eclipse Plus, 2.1 × 50 mm, 1.8 µm) with a 5 mm guard column inline. The LC method was a 0.4 mL/min gradient with water as mobile phase A and 95% methanol and 5% water as mobile phase B, both buffered with 5 mM ammonium acetate, and increased from 10 to 100% B over 16.5 minutes. The ion source was an Agilent JetStream ESI operated in negative ion mode. The IMS drift tube used a constant voltage drop and a 60 ms maximum drift time, and 4-bit multiplexing was used for the MS1-only injections. For the size-dependent AIF injections, a drift time-based collision energy ramp from 2 to 60 V was applied to alternating frames so that the larger molecules with higher drift times received a higher voltage.(*3*) The QTOF was operated in *m/z* 50-1700 mode and a mass calibration was performed every other day. MS1-only data files were demultiplexed in PNNL Preprocessor (v. 4.0)(*17*) with a signal intensity threshold of 20 counts and 100% pulse coverage. To convert measured drift times into CCS values, single-field CCS calibration was applied to all data files in Agilent IM-MS Browser (v. 10.0) using the Agilent ESI tune mix.

### Target Screening and Absolute Quantification

Peak picking was performed in Skyline-daily (v. 24.1.1)(*18, 19*) using a mass resolving power of 20,000 and a mobility resolving power of 50. Chromatograms for the internal standards were assessed to ensure retention times were within ± 0.5 min across the worklist. Samples with no internal standard signal detected were removed from the Skyline document and not included in subsequent analysis. All MS1-only files were screened using a transition list from the Baker lab LC-IMS-MS PFAS library v1, which contains retention times and CCS values for 168 PFAS standards collected under identical LC, IMS, and HRMS conditions.(*20*) Surrogate internal standards were assigned based on structural similarity and nearest retention time (**Data S4**) and used for peak area normalization (ratio of native to heavy peak area). Normalized peak areas were exported to Excel for further analysis. Detected PFAS were assigned confidence levels based on a scale adapted from existing guidance incorporating PFAS attributes and IMS-specific criteria.(*21-24*) In brief, this includes Level 1 (confirmed by reference standard analyzed under identical conditions), 2 (one probable structure supported by all available evidence), 3 (one or more tentative candidate structures), 4 (unequivocal molecular formula), and 5 (feature of interest). Calibration curves for 26 PFAS were generated in Skyline using an acceptable accuracy of 70-130%. Method reporting limit (MRLs) were defined as the average plus 3 standard deviations of the abundance in the blanks. Limits of quantitation (LOQs) were defined as the lowest point on each calibration curve that was both above the MRL and had an acceptable accuracy (**Data S6**).

### PFAS Discovery and Non-Targeted Analysis

Data files were mined for additional PFAS after completing initial target screening and absolute quantification. Agilent Mass Profiler v10.0.2.202 was used to generate a feature list from the demultiplexed MS1 files and Fluoromatch-IM v1.2 was used to score potential PFAS features using homologous series and mass defect flagging; relevant settings are provided in the **Supplementary Materials** (**Tables S5-S6**). An in-house Python-based software was used with a mass error cutoff of ± 10 ppm and a CCS error cutoff of 2.0% to reduce data by eliminating spurious detections from the feature list and to prioritize potential PFAS features which were then further examined in Agilent IM-MS Browser v10.0. Manual examination involved plotting CCS and RT vs *m/z* trendlines, searching for drift-aligned fragments in the AIF data files, and exploring literature for potential related compounds reported in water and other matrices from the same geographic region. Identified features were added to the Skyline transition list and then the target screening workflow described above was followed.

### Statistical Analysis

Statistical analyses were performed in GraphPad Prism (v10.3.1) and Python (3.12). Correlation analysis (nonparametric Spearman correlation, two-tailed p-value, 95% confidence interval) and multiple linear regression was performed to evaluate the relationship between Σ21 PFAS concentrations, sampling year, site, sex, body mass index (BMI), and life stage (adult or juvenile). Significant relationships were found between Σ21 PFAS and site (p < 0.0001), life stage (p = 0.0024), and BMI (p = 0.028), A significant interaction was also found between life stage and BMI (p < 0.0001). We elected to group adults and juveniles together despite the known interaction to ensure we had an n ≥ 3 for each site and year. A larger sample size of both age groups would be beneficial for future work; however, we note that catching even one alligator is quite challenging and sometimes you have to make the best of what you get.

## Results and Discussion

### Non-Targeted PFAS Analysis of Alligator Plasma

Plasma samples were collected from alligators in three watersheds over the 5 year study (2018-2022): the Cape Fear River downstream of a fluorochemical manufacturer in North Carolina (CFR, n = 98); Lake Waccamaw, a nearby reference site in the Lumber River basin with no known PFAS point sources (LW, n = 74); and a region in Florida with no known fluorochemical manufacturing (FL, n = 26, 2021 only) (**Figure 1, Data S1-S2**). Within the CFR, four distinct locations were sampled: Greenfield Lake (2018-2022), Orton Pond (2019 only), Oak Island (2019 only), and Bald Head Island (2019 and 2022). Data from our LC-IMS-CID-HRMS analysis was evaluated using complementary approaches combining a library screening workflow to find known PFAS (**Data S3**) and a discovery workflow to uncover additional PFAS not present in the library.

All together we identified 46 PFAS (**Table 1, Data S4-S5**) in the alligator plasma. Library screening detected 34 known PFAS (at least 45 different molecules when counting structural isomers), of which 21 were quantified (see ***Spatial and Temporal Trends*** section). The discovery workflow uncovered an additional 22 features of interest which were then manually investigated. Of these 22 features, we elucidated candidate structures for 12 PFAS (2 of which had 2 ion types detected), found fragmentation evidence for 3 PFAS with no precursor signal detected, and the remaining 5 features of interest could not be identified further. Broadly, the 12 candidate structures included 2 novel PFAS (H-substituted polyfluorinated triether acids) and 1 in-source fragment of these, 2 perfluoroalkyl ether sulfonic acids (PFAESAs) known from patent literature but not previously reported in this region, a novel isomer of PFUdA, 2 PFAS previously detected in fish (PFUdS and PFPeDA)(*19*), 1 fluorinated insecticide (fipronil sulfone), 2 new isomers of Nafion Byproduct 2 (NB2), and an unsaturated PFOS (U-PFOS, or PFOS with a double bond) isomers, which we count as 1 structure here due to our inability to determine different double bond positions. The following sections describe each structural elucidation in more detail.

**Table 1.**
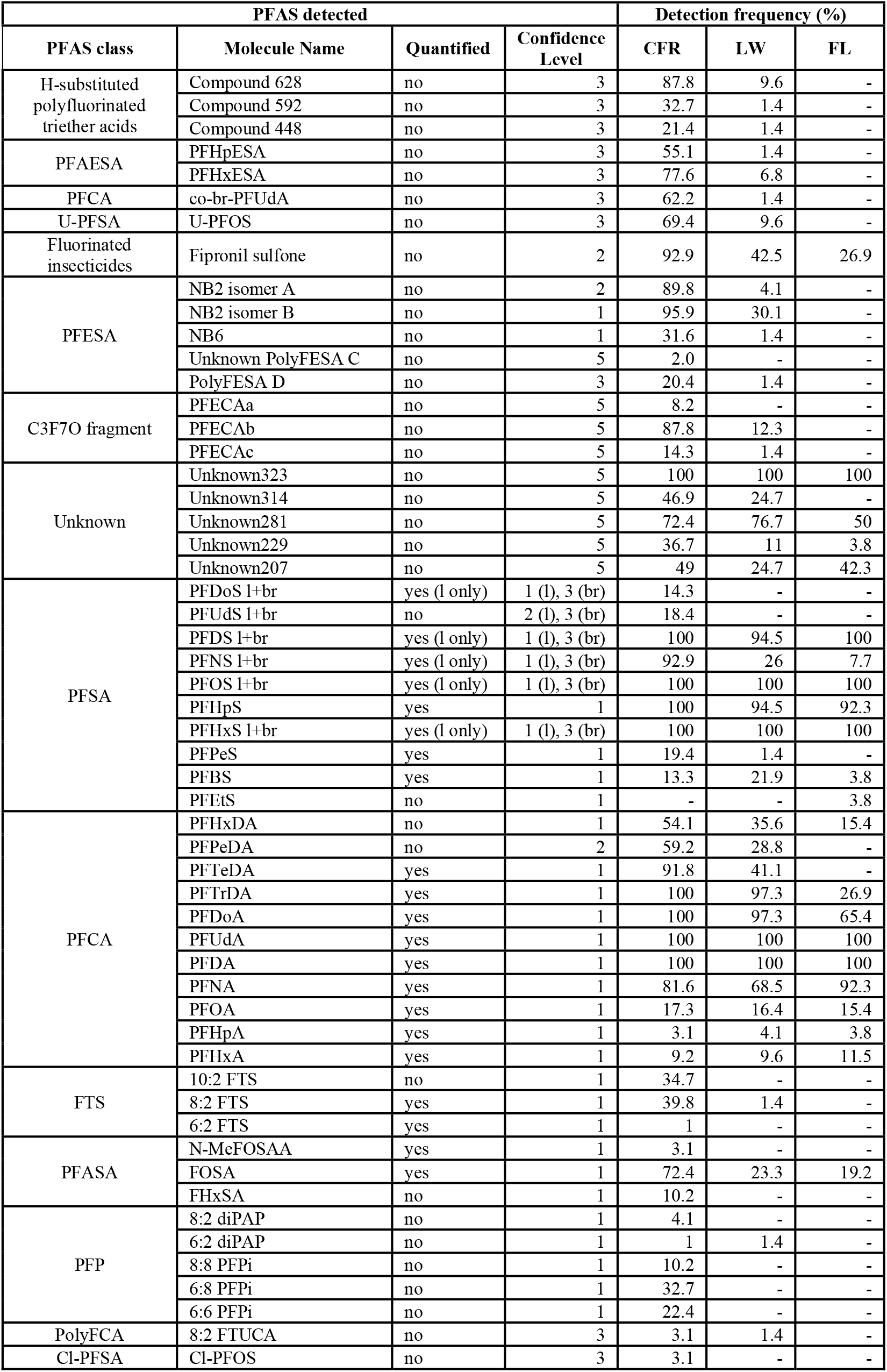
All PFAS detected in at least one alligator sample. PFAS are grouped by structural class. Detection frequency is the proportion of alligators tested at each site that had detectable levels of that molecule.

### H-substituted polyfluorinated triether acids

We report here for the first time structures for two H-substituted polyfluorinated triether acid PFAS (**Figure 2, Table 1**), referred to subsequently as Compound 628 (*m/z* 628.9180) and Compound 592 (*m/z* 592.9510). These PFAS consist of a branched fluorinated alkyl chain with one hydrogen substitution for a fluorine and three ether linkages with either a carboxylic acid or sulfonic acid headgroup. LC chromatograms for both features show several overlapping peaks at each *m/z*, indicating unresolved structural isomers or conformers. Our discovery workflow also flagged Compound 448 (*m/z* 448.9676), which was previously reported by Donelson *et al*. as an MS2 fragment of the multiheaded polyfluorinated ether Compound 10.(*3*) We did not detect Compound 10 or any of the multiheaded polyfluorinated ethers previously reported by Donelson *et al*. in the alligators. Our candidate structure for Compound 592 is desulfonated Compound 10 and our candidate for Compound 628 is decarboxylated Compound 10. Compound 448 was present in MS1-only data files along with Compound 592, followed the same LC peak shape, and had the same drift time. Together these observations suggest that it is a post-source decay product of Compound 592 (*i*.*e*. an unintentional fragment) rather than a separate feature. While fragmentation data was difficult to discern because these compounds elute closely, do not fully resolve in the drift dimension, and had generally low signal in the AIF method, predominant MS2 fragments at *m/z* 282.9822 and 262.9760 were seen in all LC peaks, indicating a loss of HF from adjacent CFH-CF_2_ or CFH-CF_3_ moieties for both features. Some of the fragments reported for Compound 10 were also seen in the alligator features; for example, Compound 592 had a distinct *m/z* 162.9824 peak (C_3_F_5_O_2_), and both features had an *m/z* 212.9792 peak (C_4_F_7_O_2_). All together this evidence suggests multiple unresolvable isomers and fragments and diagnostic evidence allowed us to propose tentative candidate structures for Compounds 592 and 628 at a Level 3 confidence(*24*) (**Figures S1-S3**).

**Figure 2.**
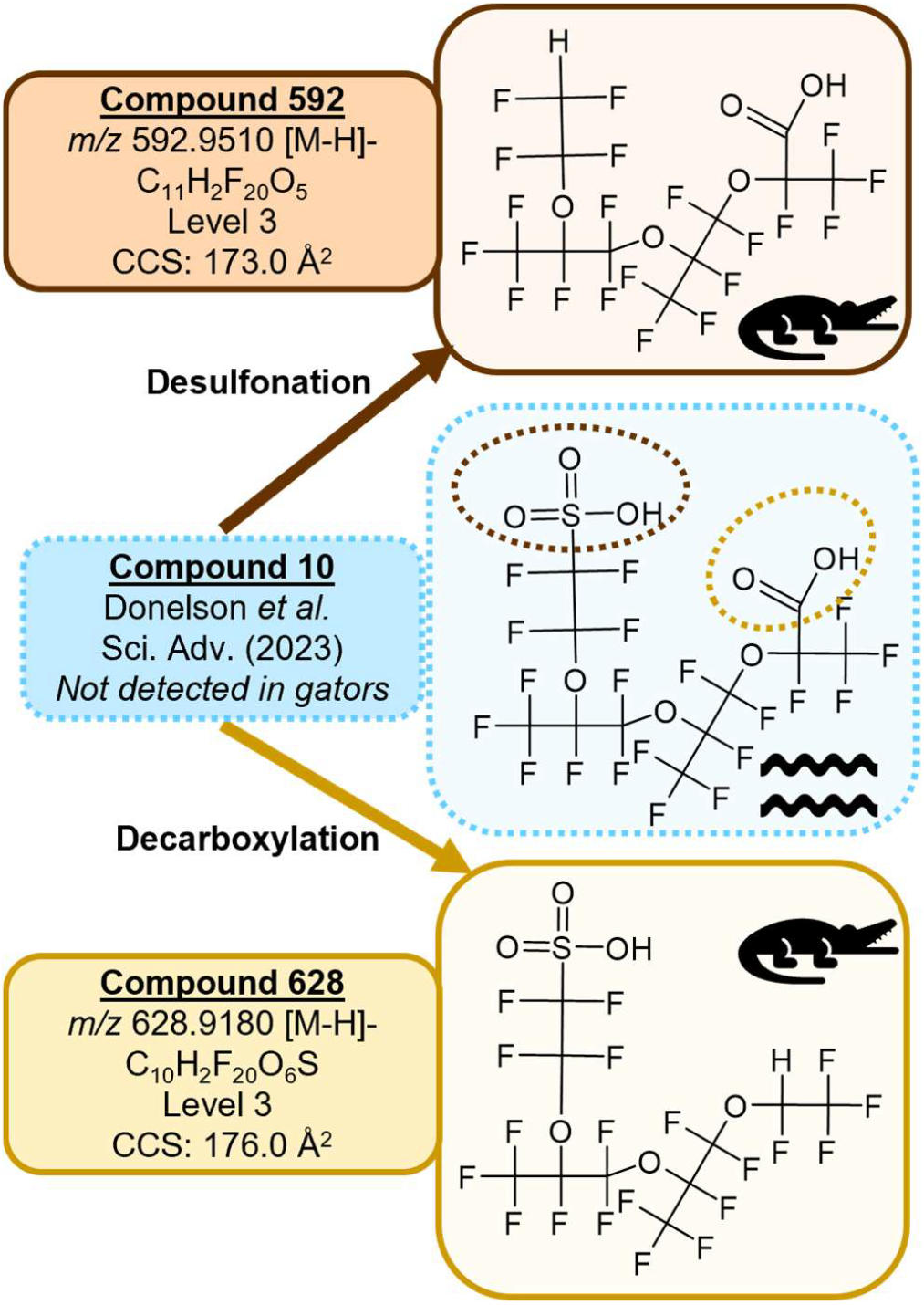
Candidate structures for the novel triether PFAS. Compounds 592 and 628 were detected in alligators and in retroactive screening of water data. Compound 10 was previously detected in water but was not detected in alligators.

Initially we hypothesized that Compounds 592 and 628 were metabolic transformation products of Compound 10. To test this, we obtained data files from Donelson *et al*. and screened CFR water samples for Compounds 592 and 628. Interestingly, both compounds were detected in the water samples, suggesting that Compounds 592 and 628 are either separate manufacturing byproducts from the Chemours facility, or possibly environmental or microbial metabolic transformation products of Compound 10. Little information is available on biological metabolism of PFAS.(*25*) Additional studies screening for all three compounds would be informative.

### Long Chain Perfluoroalkyl Ether Sulfonic Acids (PFAESAs)

Two long chain perfluoroalkyl ether sulfonic acids (PFAESAs), PFHxESA and PFHpESA, were detected (**Figure 3, Table 1**). These PFAS were reported only one time previously in biobanked European fish liver in a non-targeted study(*26*) and have never been detected in the CFR. Donelson *et al*. reported a homologous series of this subclass in passive sampling devices deployed in the CFR(*3*), including PFEESA (a known component of Nafion production(*2*)), PFPrESA, PFBESA, and PFPeESA, but we did not detect these smaller molecules in the alligator plasma. Data files were screened for a potential larger addition to the series with 8 carbons, PFOESA, but no features with an *m/z* within 10 ppm of calculated were detected. Multiple LC peaks were observed for both molecules, consistent with the peak shapes previously reported in fish liver(*26*), Trendlines using RT and CCS values from Donelson *et al*. had good fits for the most abundant PFHxESA and PFHpESA peaks, which further supported our tentative identifications. Donelson *et al*. saw fragments diagnostic of an SO_3_(CF_2_)_2_O group linked to a fluorinated alkyl tail expanding by CF_2_ units; however, we observed fragmentation patterns that suggested the presence of multiple structural isomers for PFHxESA and PFHpESA. The earlier RT peaks for PFHpESA also had smaller CCS values, which suggests more compact structural conformations that are also more polar, possibly due to branching of the fluorinated alkyl tail. However, for PFHxESA, the earlier RT peak had a larger CCS value, which suggests a more extended conformation that is more polar than the later peak, possibly due to an ether linkage closer to the CF_3_ end of the chain. Signal intensity was generally low in AIF injections, limiting our ability to propose a singular structure for each peak. However, the combination of diagnostic fragmentation evidence and CCS and RT predictions using homologous series trendlines supports presence of multiple structural isomers and allows us to identify PFHxESA and PFHpESA at Level 3 confidence(*24*) (**Figures S4-S6**).

**Figure 3.**
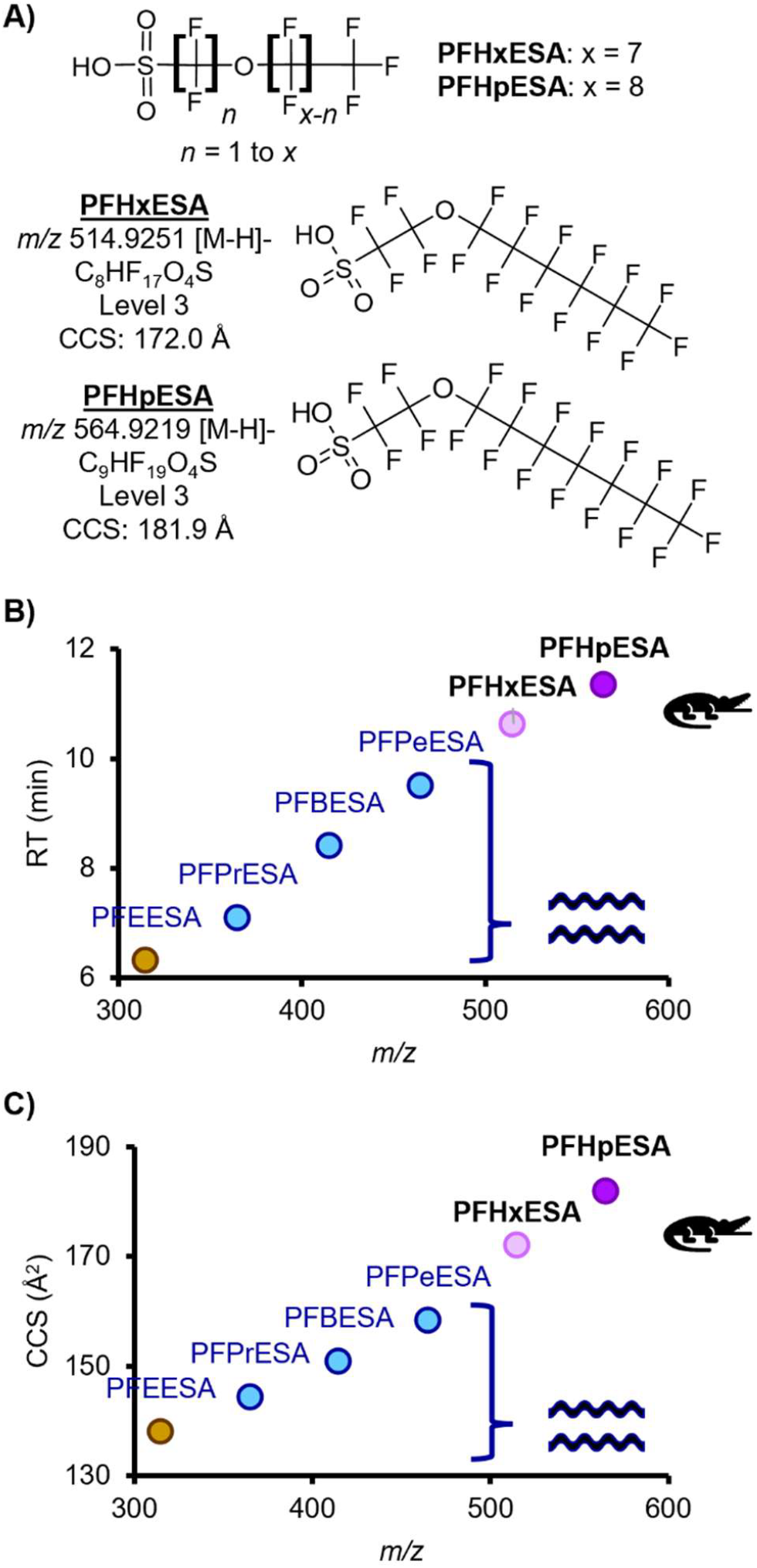
Identification of the long chain PFAESAs. (**A**) Example structure and tentative candidate structures for the two novel PFAESAs. (**B**) Retention time vs. *m/z* trendlines and (**C**) collision cross section vs *m/z* trendlines. PFEESA through PFPeESA were previously detected in water but were not detected in gators.

### Novel Perfluoroundecanoic Acid Isomer, co-br-PFUdA

A structural isomer of PFUdA, the 11-carbon carboxylic acid, was detected (**Figure 4, Table 1**). Because the proposed tentative candidate structure is branched (br) and has a non-terminal CO_2_ group, we refer to this structure as co-br-PFUdA. Fragments including C_2_F_5_, C_3_F_7_, C_4_F_9_, C_6_F_13_, and C_7_F_15_ confirmed that this feature is a structural isomer of PFUdA. Additionally, both M-H- and M-H-CO_2_-adducts were observed for co-br-PFUdA, including a second drift peak of the M-

**Figure 4.**
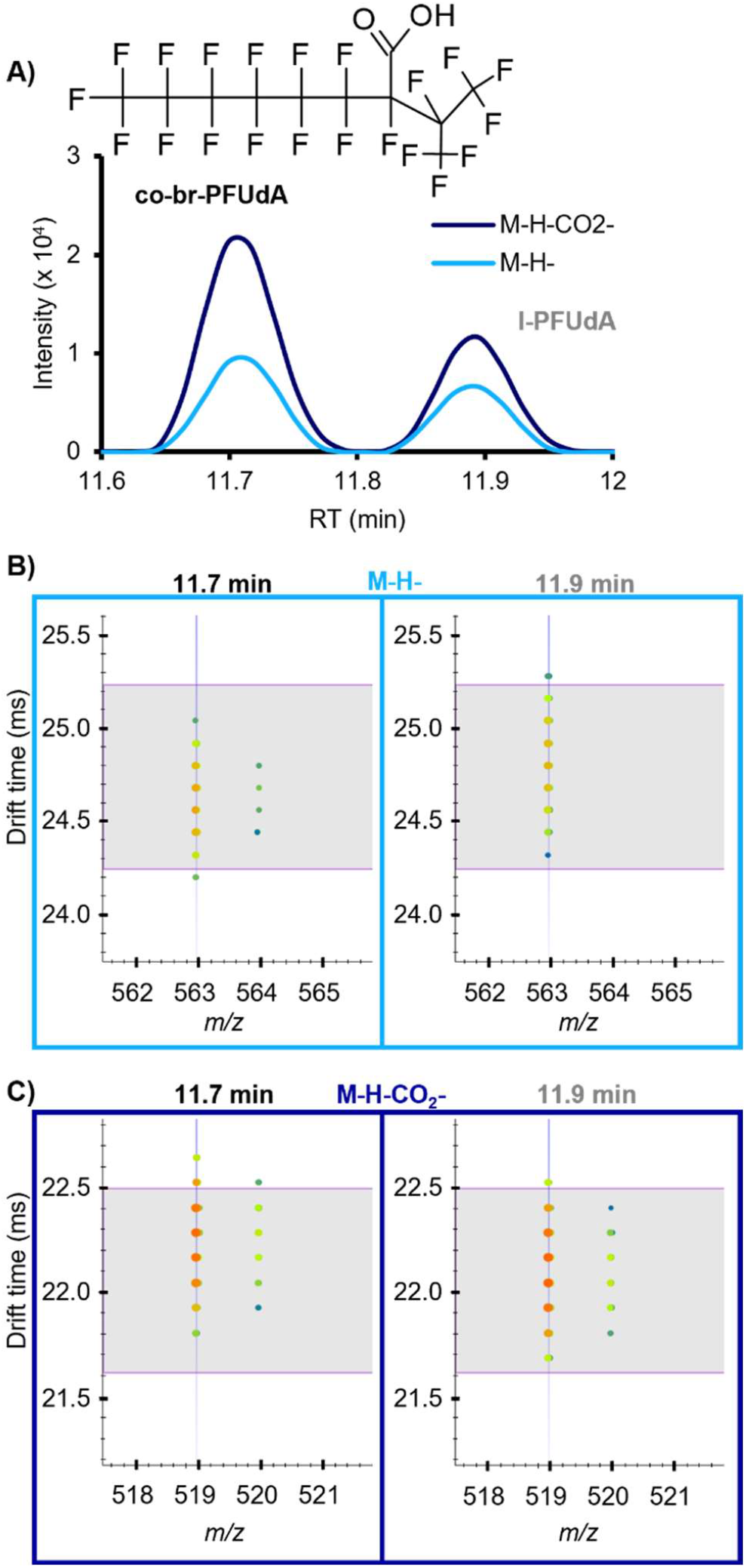
Identification of the novel PFUdA isomer. (**A**) Candidate structure of the PFUdA isomer (co-br-PFUdA) and extracted ion chromatogram illustrating the intensity of the linear PFUdA (l-PFUdA) peak at 11.9 min in comparison with the isomer at 11.7 min. (**B**) Nested drift spectra for the M-H-ions, with horizontal lines indicating the expected drift time based on the library CCS value for l-PFUdA. (**C**) Nested drift spectra for the M-H-CO_2_-ions, with horizontal lines indicating the expected drift time based on the library CCS value for l-PFUdA.

H-CO_2_-drift aligned with the M-H-, indicating post-source decay and characteristic of PFCAs on our platform. Compared with our in-house library values for l-PFUdA, the measured RT of co-br-PFUdA was slightly earlier (0.2 min) and the CCS value for co-br-PFUdA M-H-was slightly smaller (0.9%), consistent with the difference observed between other branched PFAAs and their linear counterparts. While this CCS difference is quite close, our platform is very reproducible (<0.2% error) and the RT difference allows additional confidence in the assignment. Interestingly, the M-H-CO_2_-CCS was actually 0.9% larger than that of l-PFUdA, indicating a more extended conformation upon decarboxylation and providing evidence that the CO_2_ group is in the middle of the chain rather than on the end. A PubChem formula search returned only one entry that satisfied our evidence for a CO_2_ group at a non-terminal position with a branched CF_2_ chain, which we propose as our most likely candidate structure. We can imagine several other structural possibilities for this molecule with different branching and CO_2_ locations that our evidence could support, so this identification is a Level 3 confidence (**Figures S7-S8**)(*24*). Branched isomers of carboxylic and sulfonic acid PFAS are commonly detected in lower abundance than their linear counterparts; however, this co-br-PFUdA detection is notable because its abundance was often as high or higher than the linear form, l-PFUdA, which was also detected in 100% of alligators from all sites and years.

### Other PFAS Identified Using NTA

Several other PFAS for which no standards are available were identified in our NTA workflow. We detected 3 polyfluorinated ether sulfonic acids (PolyFESAs), including 2 known isomers of Nafion Byproduct 2 (NB2 A and B, Levels 2 and 1, **Figures S9-S11**) and 1 tentative structural NB2 isomer candidate (PolyFESA D) which has not previously been reported in environmental samples (Level 3,

**Figure S13**). One additional possible PolyFESA (Unknown PolyFESA C) was originally annotated as a fourth NB2 isomer but insufficient evidence was available to confirm a tentative structure (Level 5, **Figure S12**). Additionally, we detected at least 4 other PFAS including: fipronil sulfone, a fluorinated insecticide considered to be a PFAS due to the presence of two CF_3_ groups (Level 2, **Figures S14-S15**); multiple unresolvable isomers of U-PFOS, which contain a double bond (Level 3 confidence, **Figures S16-S17**); the 15-carbon PFCA, PFPeDA (Level 2, **Figure S18**); the 11-carbon PFSA, PFUdS (Level 2, **Figure S19**). Fragmentation evidence suggested the presence of additional undetected perfluoroether carboxylic acids, but no precursor signal was detected so they could not be investigated further (**Figure S20**). Multiple U-PFOS isomers(*3*) and fipronil sulfone(*5*) were previously reported in water from the CFR. PFPeDA and PFUdS have not previously been reported in the CFR. To our knowledge, no reports of these PFAS in human residents have been made; however, their presence in alligator plasma are reason for concern.

### Spatial and Temporal Trends

While the primary focus of this work was NTA, we were also interested in evaluating how plasma concentrations of known PFAS are changing over time in these alligator populations and gathering data for comparison to other populations and future studies. Prior to sample analysis, we generated calibration curves using authenticated standards for 26 commonly targeted PFAS to allow absolute quantification (**Data S6**). Of the quantifiable PFAS, 21 of them were detected in the alligators (**Data S7-S8**). Standards do exist for the other 13 PFAS in our library that were detected and not quantified, and future work on alligators in this area should consider including them in calibration curves. Unsurprisingly, CFR alligators had the highest PFAS concentrations for the 21 quantified PFAS (denoted as Σ21) with an average of 249.1 ng/ml (median: 225.2, range: 17.3 to 806.5 ng/ml) (**Figure 5**). The LW alligators were almost 6x lower with an average of 42.1 ng/ml Σ21 PFAS (median: 33.1, range: 10.8 to 303.5 ng/ml) and FL alligators were lower still with an average of 31.8 ng/ml Σ21 PFAS (median: 30.7, range: 10.1 to 50.2 ng/ml). The composition of the Σ21 PFAS was primarily perfluoroalkyl carboxylic acids (PFCAs) and sulfonic acids (PFSAs) with median 14.8 % PFCAs and 84.9 % PFSAs in the CFR, 34.7% PFCAs and 65.3% PFSAs in LW, and 25.6% PFCAs and 74.3% PFSAs in FL. These results aligned with previous data and were expected due to the Chemours discharge in the CFR. Simple linear regression revealed no significant changes in the Σ21 PFAS concentration over time from 2018 to 2022 in either the CFR or LW alligators (LW F: 1.301, DF: 72, p-value 0.258; CFR F: 0.7962, DF: 96, p-value 0.374). Only one year of FL data was available so no time trends could be evaluated.

**Figure 5.**
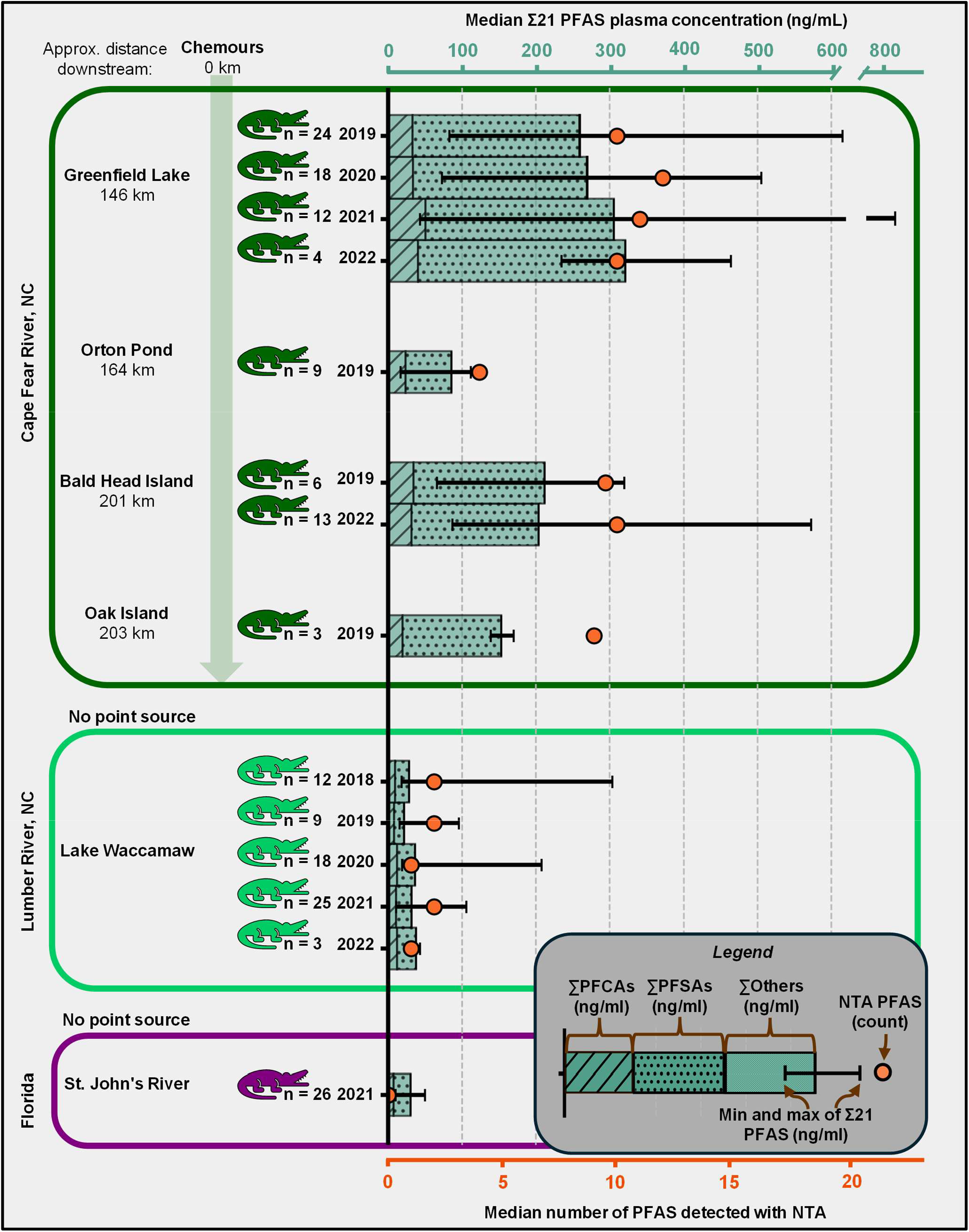
Median plasma PFAS concentrations and number of PFAS detected with NTA. The median and range of the Σ21 quantified PFAS are depicted in the bar graph (top x-axis), with the composition by subclass indicated by the colors of each bar section. Only the sites and years with n ≥3 gators sampled are shown. The orange dot indicates the number of PFAS uniquely identified using NTA (bottom x-axis), which are not included in the Σ21 PFAS concentrations.

In summary, we observed an unsurprising geographic trend with higher concentrations of Σ21 PFAS in CFR alligators compared to LW and FL alligators. No temporal trends in the Σ21 PFAS were observed at any site. Through our NTA workflow, we detected more PFAS analytes in the CFR alligators compared to LW and FL alligators, and did not observe any temporal changes in the number of PFAS detected. Importantly, the 26 PFAS identified through our NTA workflow (13 through library screening and 12 through discovery) were not quantified, and the Σ21 PFAS concentrations reported here are therefore an underestimate of the total PFAS burden in these animals. Preliminary relative abundance data for the non-quantified PFAS are provided in the **Supplementary Materials** (**Figures S21-S37** and **Data S5**). Future work will evaluate semi-quantitative methods for assessing spatial and temporal trends for the novel PFAS.

## Conclusions

Alligators live and feed exclusively in aquatic environments and represent a worst-case scenario for exposure to toxic chemicals in water. PFAS have a wide range of chemical properties and are known to partition differently between water and tissue; therefore, our objective was to see if we could detect PFAS in alligator plasma which previously had not been detected in water. Alligator plasma is a valuable biological matrix to examine for PFAS that are bioaccumulating and therefore critical to prioritize. In this study, screening against a library of CCS and RT values for over 100 PFAS allowed us to detect 34 PFAS, 21 of which were quantified. Further, we adapted an NTA workflow previously used on water from this region, which uniquely allowed us to discover 2 novel PFAS (the H-substituted polyfluorinated triether acids) and detect 3 additional PFAS (PFHxESA, PFHpESA, and co-br-PFUdA) that could not have been detected without NTA. We detected several other PFAS which have not previously been reported in biota from this region, including 3 PFAS for which standards are not available (U-PFOS, PFUdS, and PFPeDA), previously unreported isomers of Nafion Byproduct 2, and 1 fluorinated insecticide (fipronil sulfone). Detection of these compounds should be highly concerning to PFAS researchers and residents of this region because they reveal potential exposure to and uptake of dozens of compounds with unknown toxicity, in addition to very high plasma concentrations of the 21 quantified PFAS. There is an urgent need to monitor the novel and known PFAS compounds we report here so we can better understand human and wildlife exposure and increase our understanding of how dangerous these chemicals might be. Future work should assess the bioaccumulation potential of these chemicals and aim to quantify the novel compounds with no standards available. Also of note is the legality of alligator hunting in North Carolina; not only do these animals serve as useful sentinels of human exposure to PFAS, they also are a source of dietary exposure for humans who eat them.

This study illustrates a major advantage of IMS in PFAS applications, which is that all detected features have a CCS value in addition to *m/z* and RT. For homologous series, such as PFHxESA and PFHpEA, the CCS value increases with *m/z* and follows trend lines previously established for this PFAS subclass, allowing us to identify these molecules with high confidence despite their low abundance. For other novel compounds, such as Compounds 592 and 628, comparing their CCS values with *m/z* allow us to place these molecular features in the fluorinated space and enables comparison to Compound 10, a previously discovered related compound. This lends additional confidence to molecular identifications when diagnostic evidence like homologous series or manufacturing knowledge is available.(*3, 16, 27*) Several limitations are also noted for this study such as low detection limits for several classes of PFAS, such as PFECAs. A lack of internal fragmentation for some of the candidate structures also limited our ability to fully identify several of the flagged molecules. Optimization of the fragmentation method could be beneficial if full structural elucidation of the tentative candidates is required, though optimization for one class is likely to sacrifice sensitivity for other PFAS classes. Additionally, NTA requires feature prioritization that may inherently exclude certain molecules. Data analysis is highly resource intensive and time consuming, and it is not always possible to re-analyze sample extracts once a feature of interest is discovered. These challenges underscore the need for higher throughput workflows to more rapidly pull features of interest out of complex data sets, a key area of improvement needed for improved NTA.(*28*) These challenges also illustrate the benefits of IMS CCS values, which are independent of feature concentration and highly useful in molecular identifications when used in combination with additional evidence.(*3, 16, 29, 30*)

Another major advantage of NTA data collection is the potential for retroactive data mining when new analytical tools and software are developed.(*28*) We obtained data files that were collected by Donelson *et al*. from aquatic passive samplers deployed in the CFR and screened for the novel compounds detected in alligators. Notably, every novel PFAS reported here in alligators was detected retroactively in the water data. Work is ongoing to evaluate spatial and temporal trends for these novel PFAS in water. Similarly, our NTA data can be reassessed in the future as NTA tools improve. Other researchers might identify additional PFAS in our data set, which is publicly available on the MassIVE website. We urge regulators and researchers to begin monitoring the PFAS we report here in studies of food, water, wildlife, and people so we can begin to understand where these compounds come from, whether they are dangerous, and what to do about them.

## Acknowledgments

The content in this manuscript is solely the responsibility of the authors and does not necessarily reflect the views of the funding agencies. The authors wish to thank the gatoring field team for their hard work in collecting these samples. We would also like to thank the staff of North Carolina Wildlife Resource Commission and the residents of Wilmington and Lake Waccamaw, NC. AKB wishes to thank Jeff Enders for his invaluable instruction in absolute quantification and Kaylie Kirkwood-Donelson for sharing her SPATT data files for retroactive screening.

## Funding

Include all funding sources, including grant numbers, complete funding agency names, and recipient’s initials. Each funding source should be listed in a separate paragraph such as:

National Institute of Environmental Health Sciences of the National Institutes of Health Award Number P42 ES027704 (ESB)

National Institute of Environmental Health Sciences of the National Institutes of Health Award Number P42 ES031009 (SMB, ESB)

National Institute of Environmental Health Sciences of the National Institutes of Health Award Number P30 ES025128 (SMB)

National Institute of Environmental Health Sciences of the National Institutes of Health Award Number T32 ES007046 (SMB)

United States Environmental Protection Agency cooperative agreement STAR RD 84003201 (ESB)

National Science Foundation NC-C-CAPE grant OCE-2414792 (SMB)

National Institute of Environmental Health Sciences NC-C-CAPE grant P01 ES035542 (SMB)

## Author contributions

Conceptualization: AKB, SMB, ESB

Methodology: AKB

Software: AKB, GPK

Investigation: AKB, KDR

Resources: MPG, FR, SMB, ESB

Visualization: AKB, GPK

Supervision: ESB

Writing—original draft: AKB

Writing—review & editing: AKB, GPK, KDR, SMB, ESB

## Competing interests

Authors declare that they have no competing interests.

## Data and materials availability

All data are available in the main text or the supplementary materials. Skyline files containing LC-IMS-HRMS data are available at <https://panoramaweb.org/2025-alligator-PFAS.url> and raw data files are available at <ftp://massive.ucsd.edu/v09/MSV000097330/>.

## Supplementary Materials

Supplementary Materials (PDF)

Supplementary Data (XLSX)

## Notes

https://panoramaweb.org/2025-alligator-PFAS.url

ftp://massive.ucsd.edu/v09/MSV000097330/

### Competing Interest Statement

The authors have declared no competing interest.

### Summary of Updates

This version of the manuscript was revised to address reviewer comments.

## References

1. M. Sun et al., Legacy and emerging perfluoroalkyl substances are important drinking water contaminants in the Cape Fear River Watershed of North Carolina. Environmental science & technology letters 3, 415–419 (2016).

2. J. McCord, M. Strynar, Identification of Per-and Polyfluoroalkyl Substances in the Cape Fear River by High Resolution Mass Spectrometry and Nontargeted Screening. Environ Sci Technol 53, 4717–4727 (2019).

3. K. I. Kirkwood-Donelson, J. N. Dodds, A. Schnetzer, N. Hall, E. S. Baker, Uncovering per- and polyfluoroalkyl substances (PFAS) with nontargeted ion mobility spectrometrymass spectrometry analyses. Sci Adv 9, eadj7048 (2023).

4. N. Kotlarz et al., Per- and polyfluoroalkyl ether acids in well water and blood serum from private well users residing by a fluorochemical facility near Fayetteville, North Carolina. J Expo Sci Environ Epidemiol 34, 97–107 (2024).

5. R. A. Weed et al., Non-Targeted PFAS Suspect Screening and Quantification of Drinking Water Samples Collected through Community Engaged Research in North Carolina’s Cape Fear River Basin. Toxics 12, (2024).

6. A. R. Robuck et al., Tissue-specific distribution of legacy and novel per- and polyfluoroalkyl substances in juvenile seabirds. Environ Sci Technol Lett 8, 457–462 (2021).

7. H. M. Pickard et al., PFAS and Precursor Bioaccumulation in Freshwater Recreational Fish: Implications for Fish Advisories. Environmental Science & Technology 56, 15573–15583 (2022).

8. A. O. De Silva et al., PFAS Exposure Pathways for Humans and Wildlife: A Synthesis of Current Knowledge and Key Gaps in Understanding. Environmental Toxicology and Chemistry 40, 631–657 (2021).

9. J. Bangma et al., Combined screening and retroactive data mining for emerging perfluoroethers in wildlife and pets in the Cape Fear region of North Carolina. Chemosphere 363, 142898 (2024).

10. PFAS Testing Results (2017 https://www.cfpua.org/833/PFAS-Testing-Results).

11. T. C. Guillette et al., Elevated levels of per- and polyfluoroalkyl substances in Cape Fear River Striped Bass (Morone saxatilis) are associated with biomarkers of altered immune and liver function. Environment International 136, 105358 (2020).

12. N. Kotlarz et al., Measurement of Novel, Drinking Water-Associated PFAS in Blood from Adults and Children in Wilmington, North Carolina. Environ Health Perspect 128, 77005 (2020).

13. K. D. Rock et al., Domestic Dogs and Horses as Sentinels of Per- and Polyfluoroalkyl Substance Exposure and Associated Health Biomarkers in Gray’s Creek North Carolina. Environmental Science & Technology 57, 9567–9579 (2023).

14. S. M. Belcher, M. P. Guillette, F. Robb, K. D. Rock, Comparative assessment of blood mercury in American alligators (Alligator mississippiensis) from Coastal North Carolina and Florida. Ecotoxicology 31, 1137–1146 (2022).

15. T. C. Guillette, T. W. Jackson, M. Guillette, J. McCord, S. M. Belcher, Blood concentrations of per- and polyfluoroalkyl substances are associated with autoimmune-like effects in American alligators from Wilmington, North Carolina. Front Toxicol 4, 1010185 (2022).

16. J. N. Dodds, Z. R. Hopkins, D. R. U. Knappe, E. S. Baker, Rapid Characterization of Per- and Polyfluoroalkyl Substances (PFAS) by Ion Mobility Spectrometry-Mass Spectrometry (IMS-MS). Anal Chem 92, 4427–4435 (2020).

17. A. Bilbao et al., A Preprocessing Tool for Enhanced Ion Mobility-Mass Spectrometry-Based Omics Workflows. J Proteome Res 21, 798–807 (2022).

18. K. I. Kirkwood et al., Utilizing Skyline to analyze lipidomics data containing liquid chromatography, ion mobility spectrometry and mass spectrometry dimensions. Nature Protocols 17, 2415–2430 (2022).

19. A. K. Boatman et al., Assessing Per- and Polyfluoroalkyl Substances in Fish Fillet Using Non-Targeted Analyses. Environmental Science & Technology 58, 14486–14495 (2024).

20. K. Joseph, Boatman, A., Dodds, J., Kirkwood Donelson, K., Ryan, J., Zhang, J., Thiessen, P., Bolton, E., Valdiviezo, A., Sapozhnikova, Y., Rusyn, I., Schymanski, E., & Baker, E.. (Zenodo, 2024).

21. E. L. Schymanski et al., Identifying small molecules via high resolution mass spectrometry: communicating confidence. Environ Sci Technol 48, 2097–2098 (2014).

22. A. Celma et al., Improving Target and Suspect Screening High-Resolution Mass Spectrometry Workflows in Environmental Analysis by Ion Mobility Separation. Environ Sci Technol 54, 15120–15131 (2020).

23. J. A. Charbonnet et al., Communicating Confidence of Per- and Polyfluoroalkyl Substance Identification via High-Resolution Mass Spectrometry. Environ Sci Technol Lett 9, 473–481 (2022).

24. A. K. Boatman et al., Updated Guidance for Communicating PFAS Identification Confidence with Ion Mobility Spectrometry. bioRxiv, 2025.2001.2027.634925 (2025).

25. D. A. Dukes, C. A. McDonough, N-glucuronidation and excretion of perfluoroalkyl sulfonamides in mice following ingestion of aqueous film-forming foam. Environmental Toxicology and Chemistry, (2024).

26. S. Dudasova et al., An automated and high-throughput data processing workflow for PFAS identification in biota by direct infusion ultra-high resolution mass spectrometry. Anal Bioanal Chem 416, 4833–4848 (2024).

27. J. N. Dodds et al., From Pesticides to Per- and Polyfluoroalkyl Substances: An Evaluation of Recent Targeted and Untargeted Mass Spectrometry Methods for Xenobiotics. Anal Chem 93, 641–656 (2021).

28. M. Strynar et al., Practical application guide for the discovery of novel PFAS in environmental samples using high resolution mass spectrometry. J Expo Sci Environ Epidemiol 33, 575–588 (2023).

29. M. Foster et al., Uncovering PFAS and Other Xenobiotics in the Dark Metabolome Using Ion Mobility Spectrometry, Mass Defect Analysis, and Machine Learning. Environ Sci Technol 56, 9133–9143 (2022).

30. K. E. Burnum-Johnson et al., Ion Mobility Spectrometry and the Omics: Distinguishing Isomers, Molecular Classes and Contaminant Ions in Complex Samples. Trends Analyt Chem 116, 292–299 (2019).

