## Supplementary Material for "Novel PFAS in Alligator Blood Discovered with Non-Targeted Ion Mobility Spectrometry-Mass Spectrometry"

Anna Boatman *et al.*

**This PDF file includes:**

Supplementary Text

Tables S1 to S6

Figs. S1 to S37

**Other Supplementary Materials for this manuscript include the following:**

Data S1 to S8

Supplementary Text

Plasma Sample Preparation

Three days prior to extraction, 50 µl plasma was aliquoted from original collected samples into 1.5 ml polypropylene tubes and stored in the refrigerator. Pooled samples were created by combining equal volume aliquots of plasma from all animals collected at each site in each season; 50 µl aliquots of each pool were collected and extracted. Multiple 50 µl aliquots of field blank water and previously-reconstituted NIST SRM 1957 (referred to as “SRM” hereafter) were also transferred and stored under the same conditions for use as QC samples. Aliquots were blocked by site and year and then randomized into four extraction batches to limited capacity in the speedvac. Three method blanks and three SRM samples were extracted with each batch to account for background contamination and reproducibility between batches. All four batches were prepped and extracted on the same day using the same solvents. The pooled samples and calibration curve samples were prepared in a separate fifth batch on the following day using the same internal standards and extraction solvents as had been used for the individual unknown samples.

Internal standards (MPFAC-HIF-ES diluted 1:1 in methanol, 5 µl) were spiked into each plasma, field blank, or SRM aliquot. One analyst performed all of these spikes to ensure maximal reproducibility between samples and batches. Samples were vortexed briefly. 300 µl of acetonitrile (chilled to -20 °C in the freezer ahead of time) was added to each sample using the repeater pipette. Samples were again vortexed and then placed in the freezer (-20 °C) for 30 minutes. Again, one analyst performed the acetonitrile addition for all samples to minimize variation. Samples were again vortexed and then centrifuged at 12,500g and 4 °C for 5 minutes to collect precipitate. 200 µl of the acetonitrile top layer was carefully transferred into new 1.5 ml polypropylene microcentrifuge tubes, all by one analyst. Samples were evaporated to dryness in the speedvac at 45 °C and full vacuum. All batches dried in under one hour. Samples were reconstituted by adding 100 µl of 40:60 methanol:water buffered with 3 mM ammonium acetate and vortexing. Minimal precipitate was observed in the evaporated samples so the entire volume was transferred into polypropylene LC vial inserts (Agilent). Samples were stored in the refrigerator until analysis.

The 8-point matrix-matched calibration curve (7 concentrations of native standards plus a zero) was prepared using charcoal-stripped fetal bovine serum following the same extraction procedure, but with additional native standards spiked in (HFPO-DA and PFAC-MXH from Wellington; 7 different concentrations ranging from neat standard to a 31,250 fold dilution in methanol, plus a zero point of methanol only). All calibration curve samples also had one concentration of heavy standards spiked in to match the internal standards used in the unknown samples (5 µl of MPFAC-HIF-ES diluted 1:1 in methanol). 8 aliquots of 50 µl of charcoal-stripped fetal bovine serum were transferred into 2 ml polypropylene tubes. 3 x 50 µl water was prepared for method blanks, and 3 x 50 µl NIST SRM 1957 was prepared for validation samples. 5 µl mass-labeled internal standard was spiked into each sample first, followed by 20 µl of each MXH dilution (or neat methanol for the zero point) and 4 µl of each HFPO-DA dilution. Samples were then vortexed and the same extraction procedure detailed above was followed, however, only 276 µl of acetonitrile was added to each sample so that total sample volumes would be identical to the unknown samples.

A double blank of reconstitution buffer was added directly to a polypropylene LC vial. A standard blank (“SIL”) was also created by adding 10 µl of internal standard to 190 µl of recon buffer.

LC-IMS-CID-HRMS Data Collection

All samples were first analyzed in MS1-only mode (no collision energy). The pooled samples and several representative individual alligator samples were reinjected using a size-dependent all-ions fragmentation method. The LC method was a 0.4 mL/min gradient with water as mobile phase A and 95% methanol and 5% water as mobile phase B, both buffered with 5 mM ammonium acetate, and increased from 10 to 100% B over 16.5 minutes (Table S1). The ion source was an Agilent JetStream ESI operated in negative ion mode (Table S2). The IMS drift tube used a constant voltage drop and a 60 ms maximum drift time (Table S3) The QTOF was operated in m/z 50-1700 mode and 4-bit multiplexing was used for the MS1-only injections (Table SM3). For the size-dependent AIF injections, a drift time-based collision energy ramp from 2 to 60 V (Table S4) was applied to alternating frames so that the larger molecules with higher drift times received a higher voltage.

| **Time (min)** | **% B** |
| --- | --- |
| 0 | 10 |
| 0.5 | 10 |
| 2 | 30 |
| 14 | 95 |
| 16.5 (stop time) | 100 |
| 6 (post time) | 10 |

Table S1.

LC gradient settings.

| **Parameter** | **Value** | **Units** |
| --- | --- | --- |
| Gas temperature | 230 | °C |
| Drying gas flow | 11 | L/min |
| Nebulizer gas pressure | 45 | psig |
| Sheath gas temperature | 350 | °C |
| Sheath gas flow | 11 | L/min |
| Vcap | 3500 | V |
| Nozzle voltage | 500 | V |

Table S2.

Electrospray ionization source settings.

| **Parameter** | **Value** | **Units** |
| --- | --- | --- |
| Mass range | 50-1700 | *m/z* |
| Trap fill time | 3900 | µs |
| Trap release time | 300 | µs |
| Frame rate | 1 | frames/sec |
| IM transient rate | 17 | transients/frame |
| Max drift time | 60 | ms |
| TOF transient rate | 496 | transients/IM transient |
| Multiplexing pulse sequence length | 4 | bit |
| Drift tube entrance voltage | -1574 | V |
| Drift tube exit voltage | -224 | V |
| Rear funnel entrance voltage | -217.5 | V |
| Rear funnel exit voltage | -45 | V |

Table S3.

IMS-MS settings.

| **Drift time (ms)** | **Collision Energy (V)** |
| --- | --- |
| 0 | 2 |
| 10 | 5 |
| 15 | 19 |
| 21 | 32 |
| 25 | 41 |
| 35 | 53 |
| 45 | 60 |
| 60 | 60 |

Table S4.

Collision energy ramp settings.

| **Parameter** | **Value** |
| --- | --- |
| Ion intensity | >= 0.0 counts |
| Isotope model | Common organic molecules |
| Charge state | z = 1 |
| RT tolerance | ± 0.0% + 1.00 min |
| DT tolerance | ± 1.5% |
| Mass tolerance | ± 10.0 ppm + 2.0 mDa |
| Sample occurrence: Frequency | >= 10.0% across all samples |
| Group difference: Expression | Fold change >= 1.0 |

Table S5.

Mass Profiler settings.

| **Parameter** | **Value** |
| --- | --- |
| Mass window | 0.01 Da |
| CCS window | 2% |
| Mass defect flag: upper | 0.12 |
| Mass defect flag: lower | -0.11 |
| Polarity | Neg |

Table S6.

Fluoromatch-IM settings.

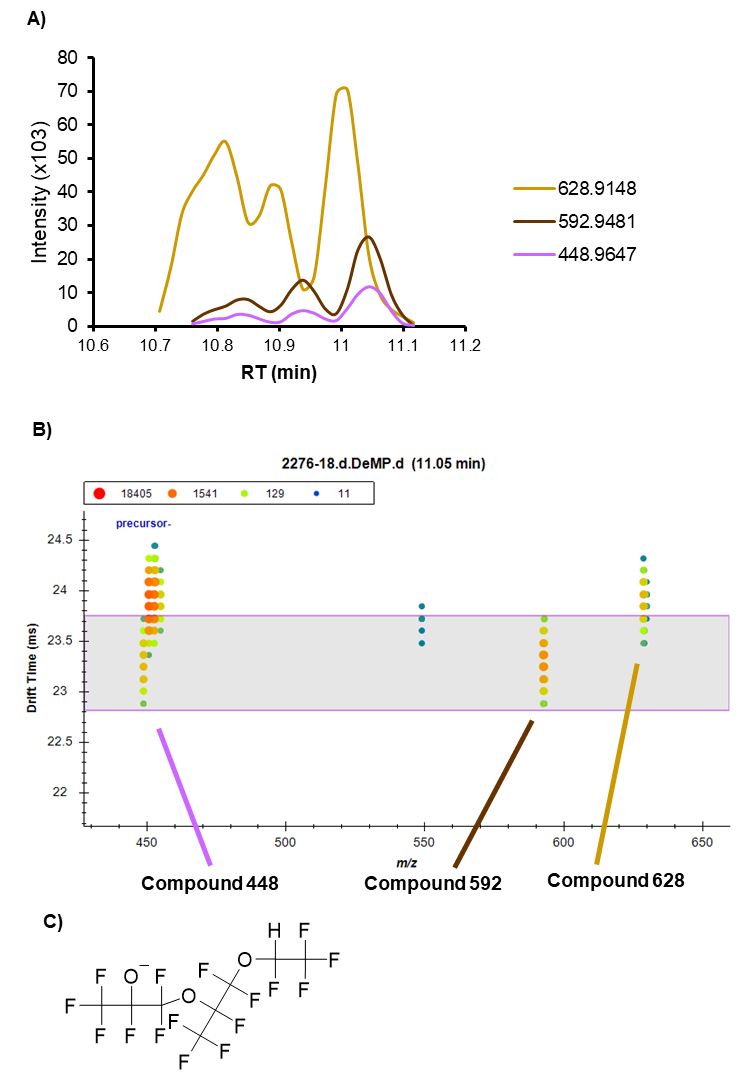

Fig. S1.

MS1-only data supporting novel triether compounds. A) Extracted ion chromatogram (EIC) of a representative sample. B) Nested drift spectra for the predominant LC peak of all 3 compounds. C) Candidate structure for Compound 448. Together this supports Compound 448 as an in source fragment of Compound 592.

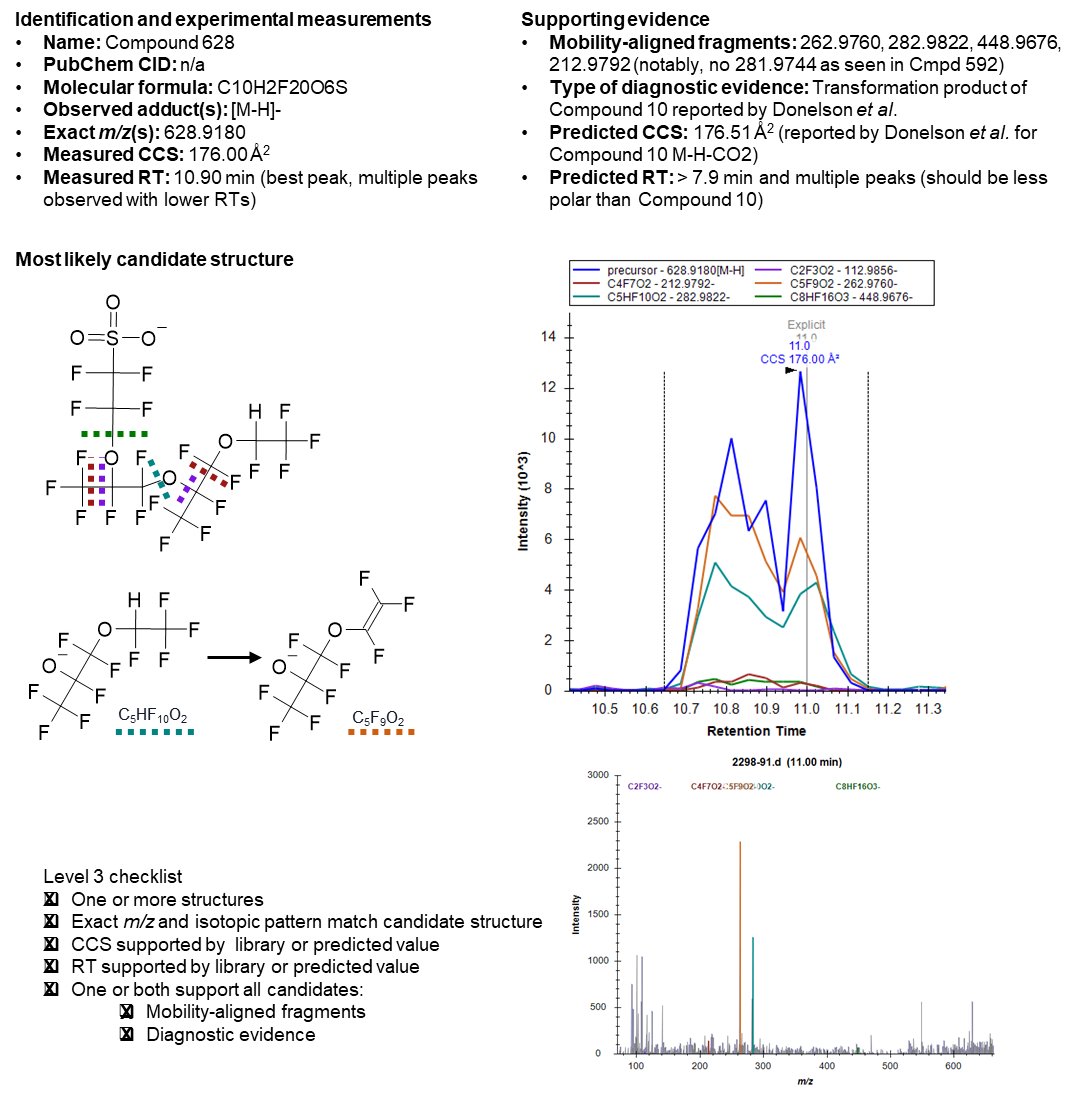

Fig. S2.

Supporting evidence for Compound 628.

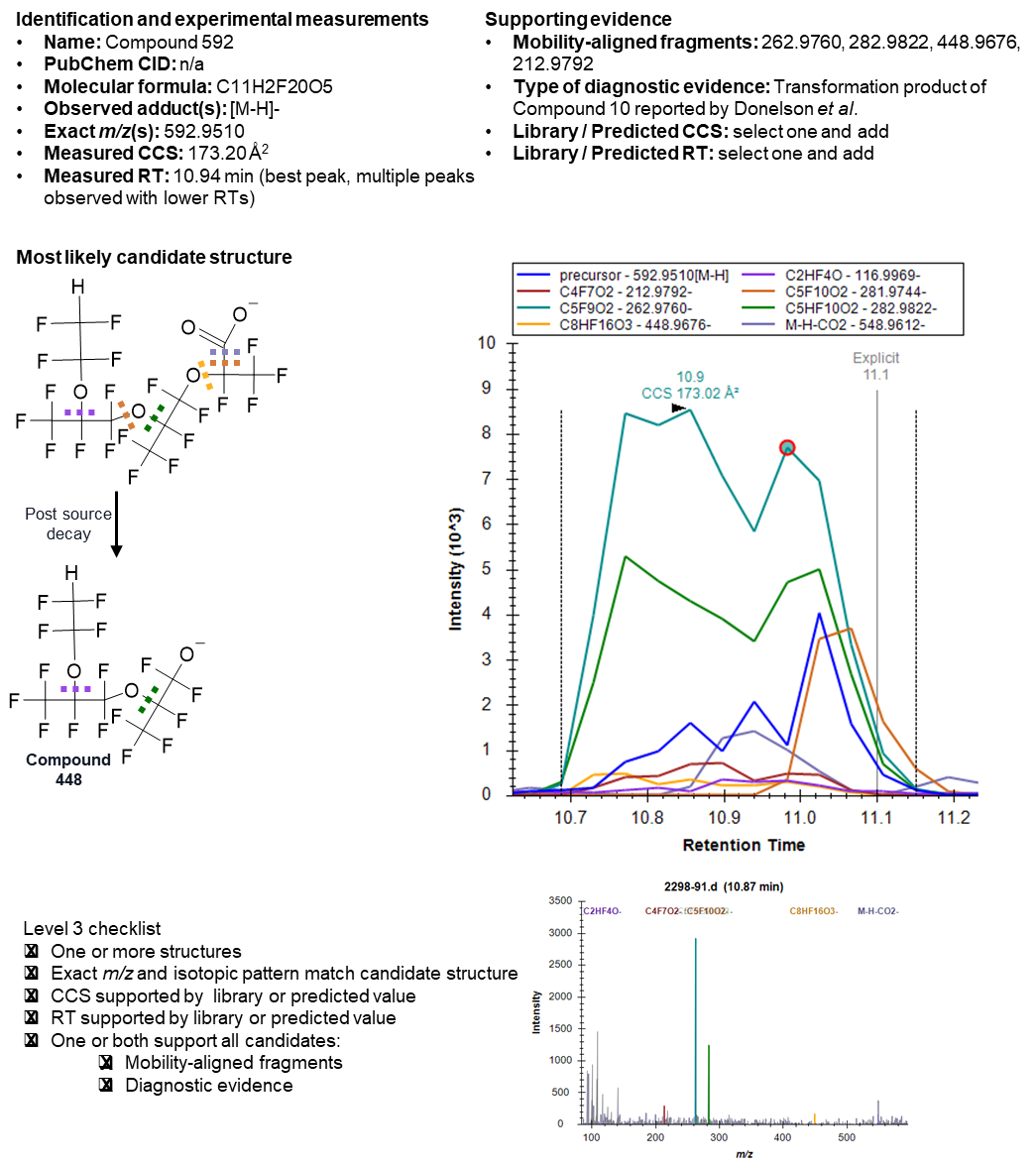

Fig. S3.

Supporting evidence for Compound 592.

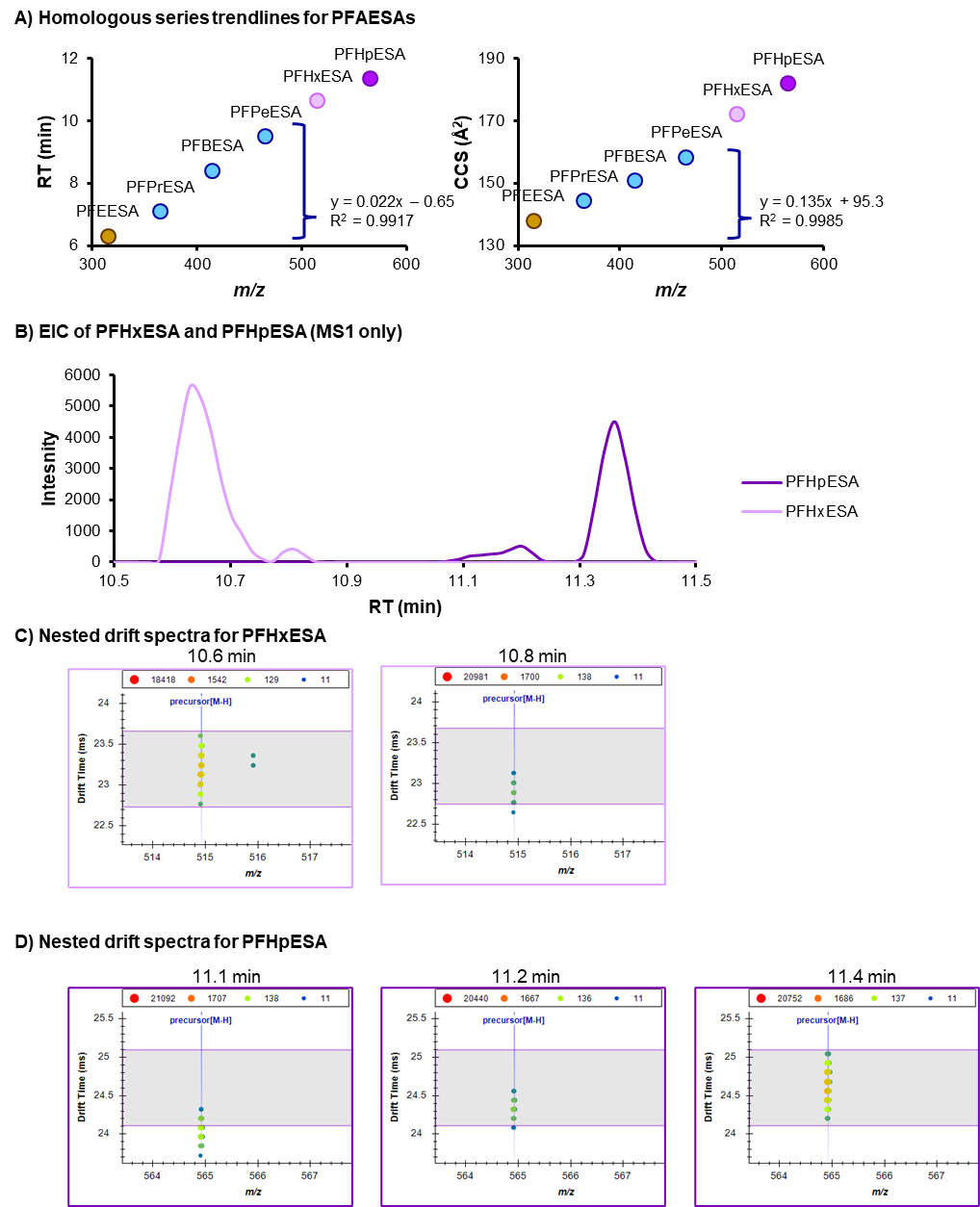

Fig. S4.

MS1-only data supporting novel PFAESAs. A) RT and CCS vs m/z trendlines for the homologous series using library values for the smaller chain molecules. B) Extracted ion chromatogram (EIC) from a representative sample showing the multiple peaks detected. C) Nested drift spectra for PFHxESA. The lower CCS at the higher RT indicates that the ether linkage is different between the two isomers. A smaller CCS indicates a more bent or compact structure while a higher RT is indicative of a longer fluoroalkyl tail, which would have more interactions with the C18 stationary phase compared to a tail with an ether linkage closer to the terminus. D) Nested drift spectra for PFHpESA.

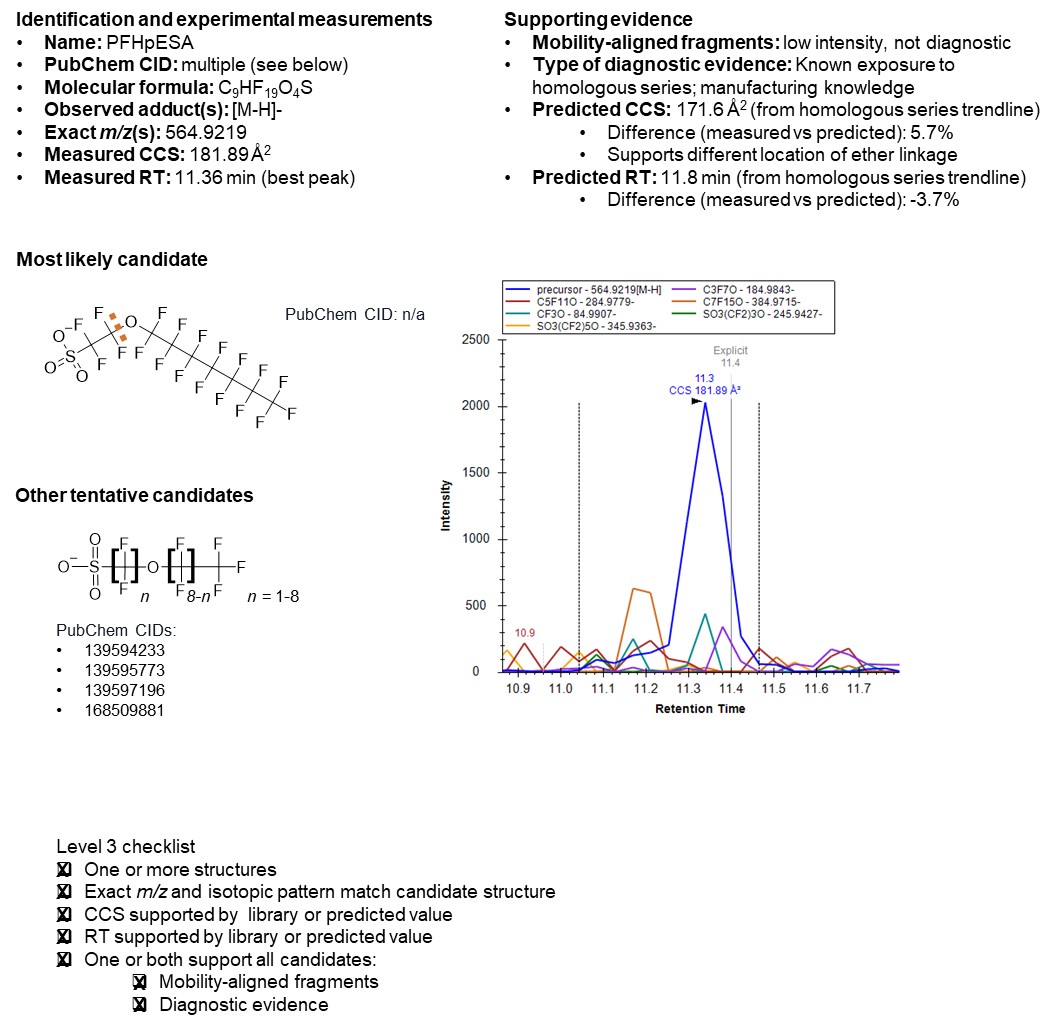

Fig. S5.

Supporting evidence for PFHpESA.

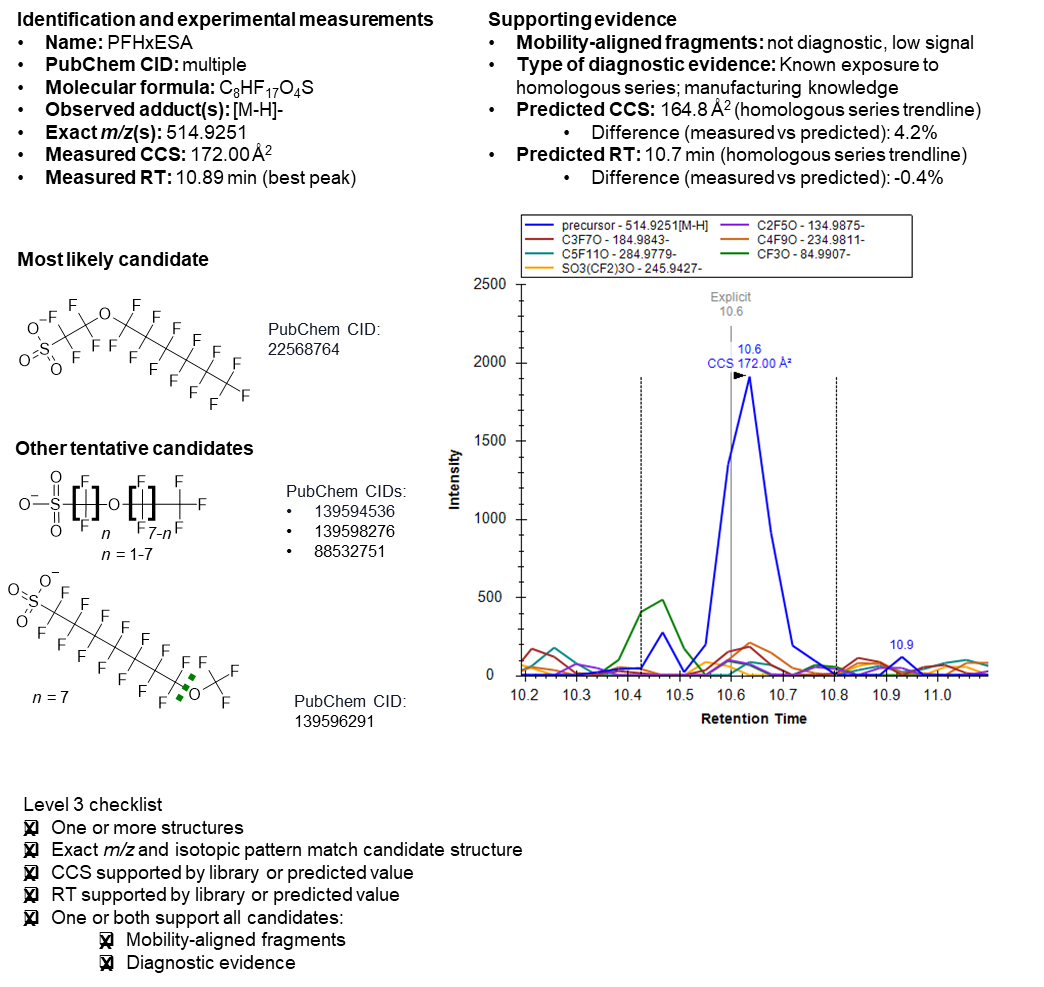

Fig. S6.

Supporting evidence for PFHxESA.

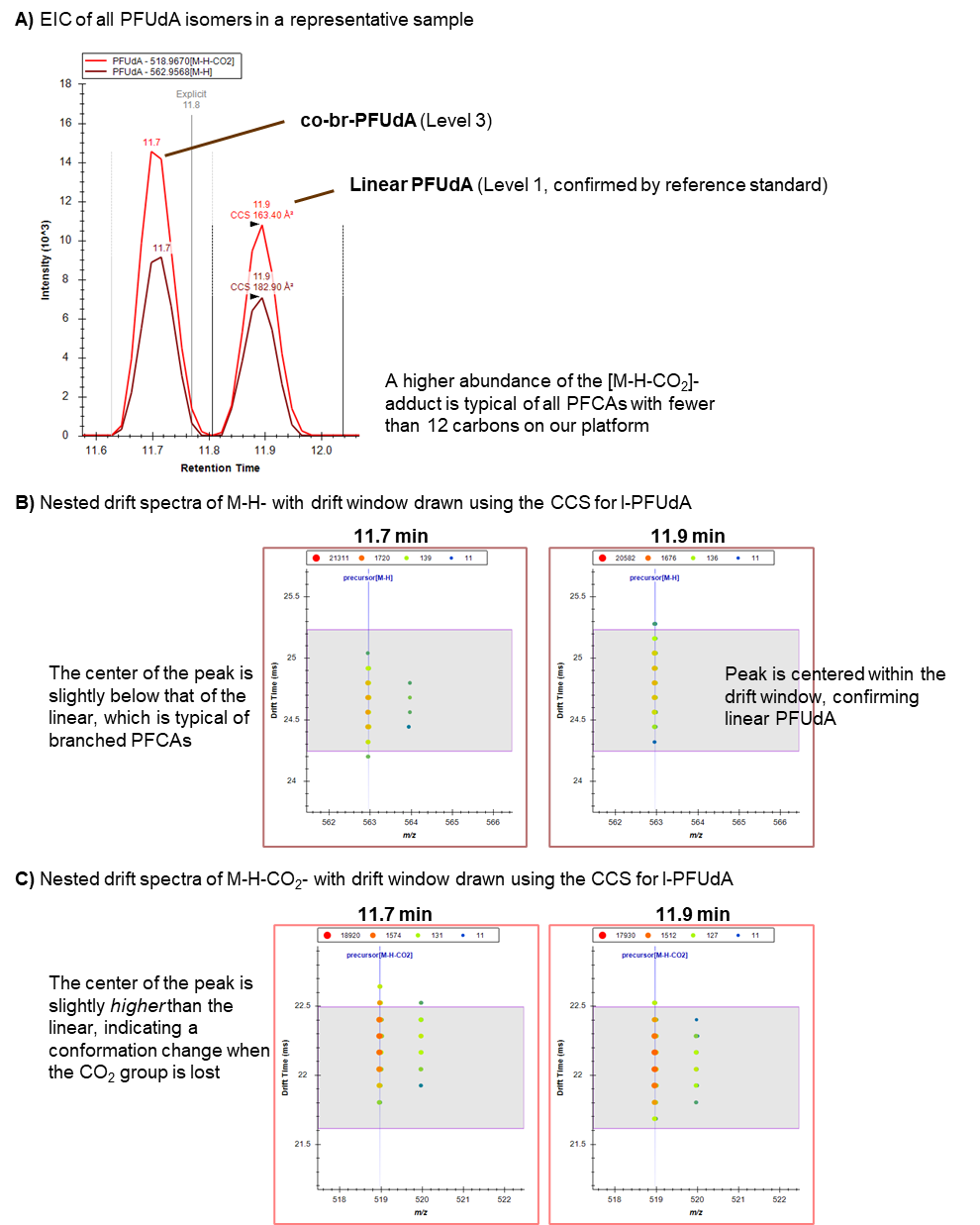

Fig. S7.

MS1-only data supporting co-br-PFUdA. A) Extracted ion chromatogram (EIC) from a representative sample showing the M-H- and M-H-CO2- ions detected for both linear PFUdA and the isomer. B) Nested drift spectra for the M-H- ion. C) Nested drift spectra for the M-H-CO2- ion.

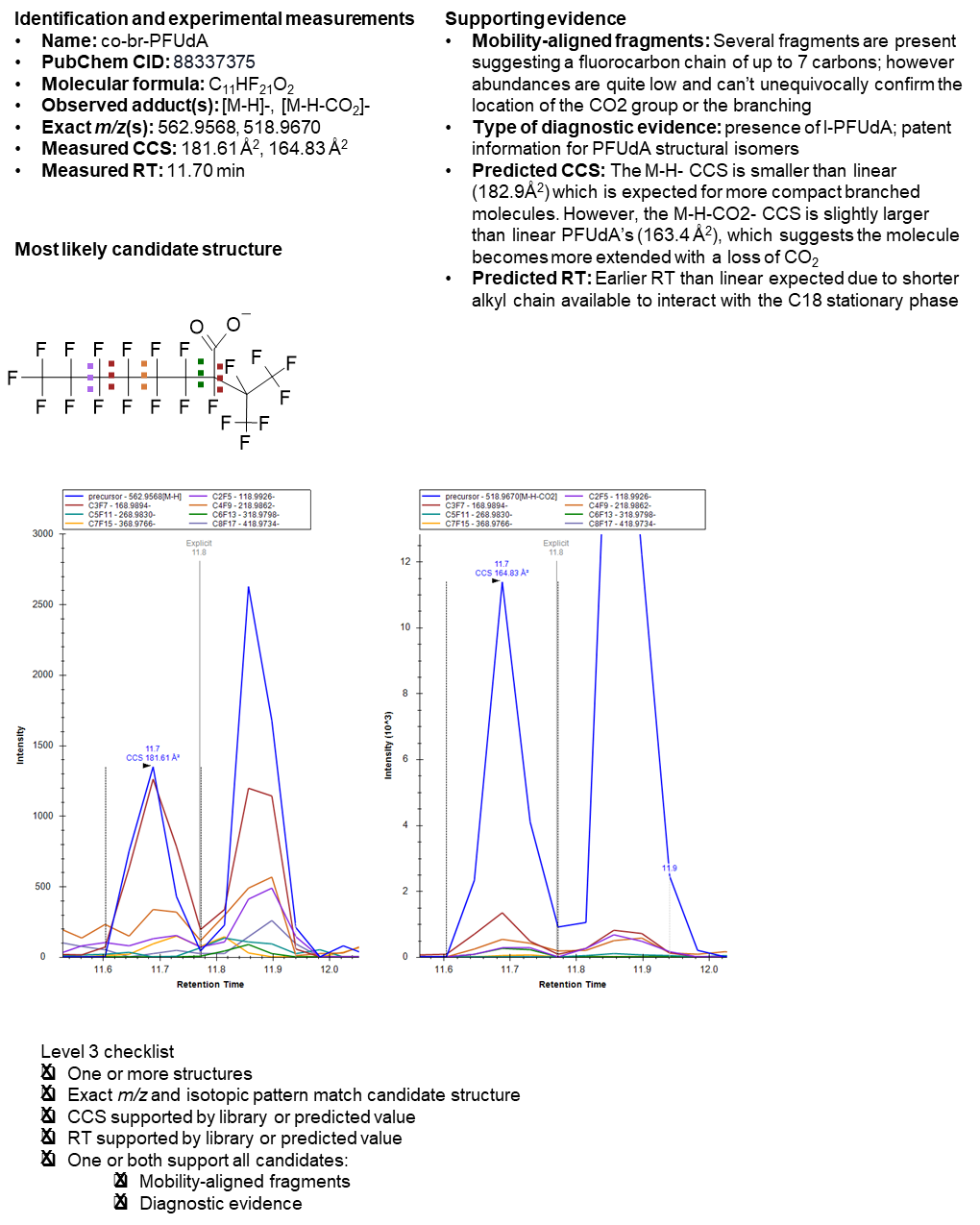

Fig. S8.

Supporting evidence for co-br-PFUdA.

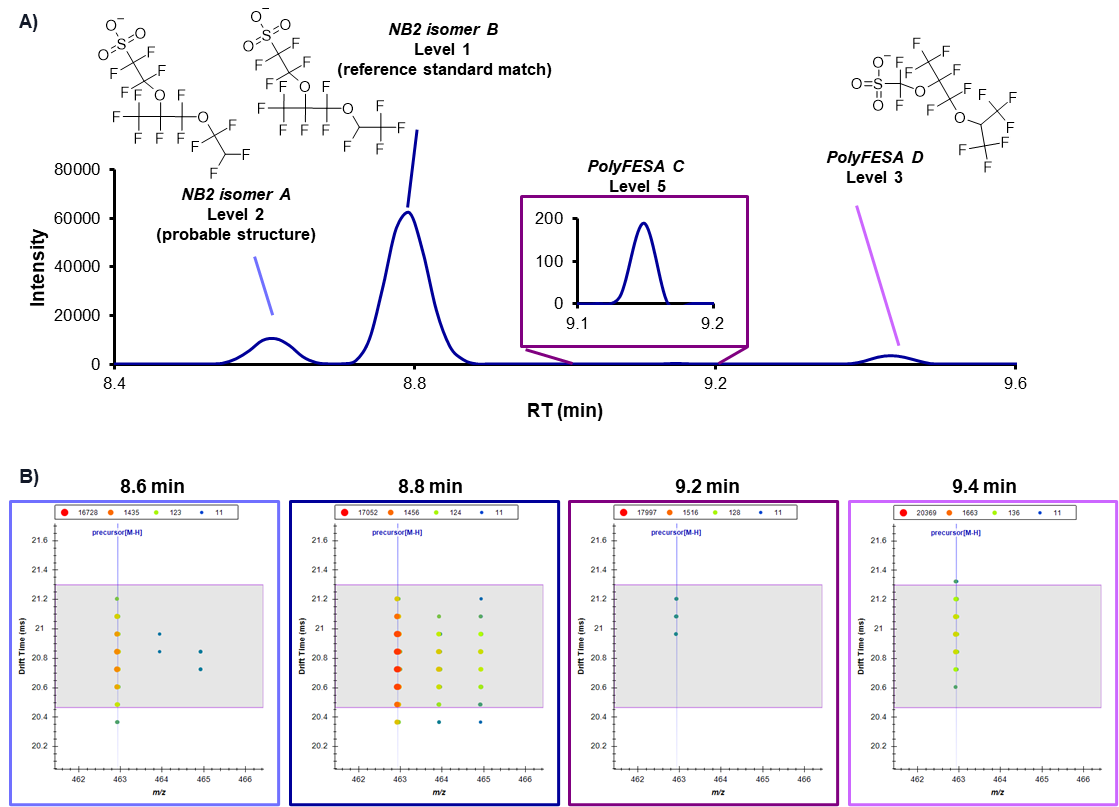

Fig. S9.

A) Extracted ion chromatogram (EIC) from a representative sample showing peaks for all 4 detected isomers, including two known Nafion Byproduct 2 (NB2) isomers. B) Nested drift spectra for each LC peak.

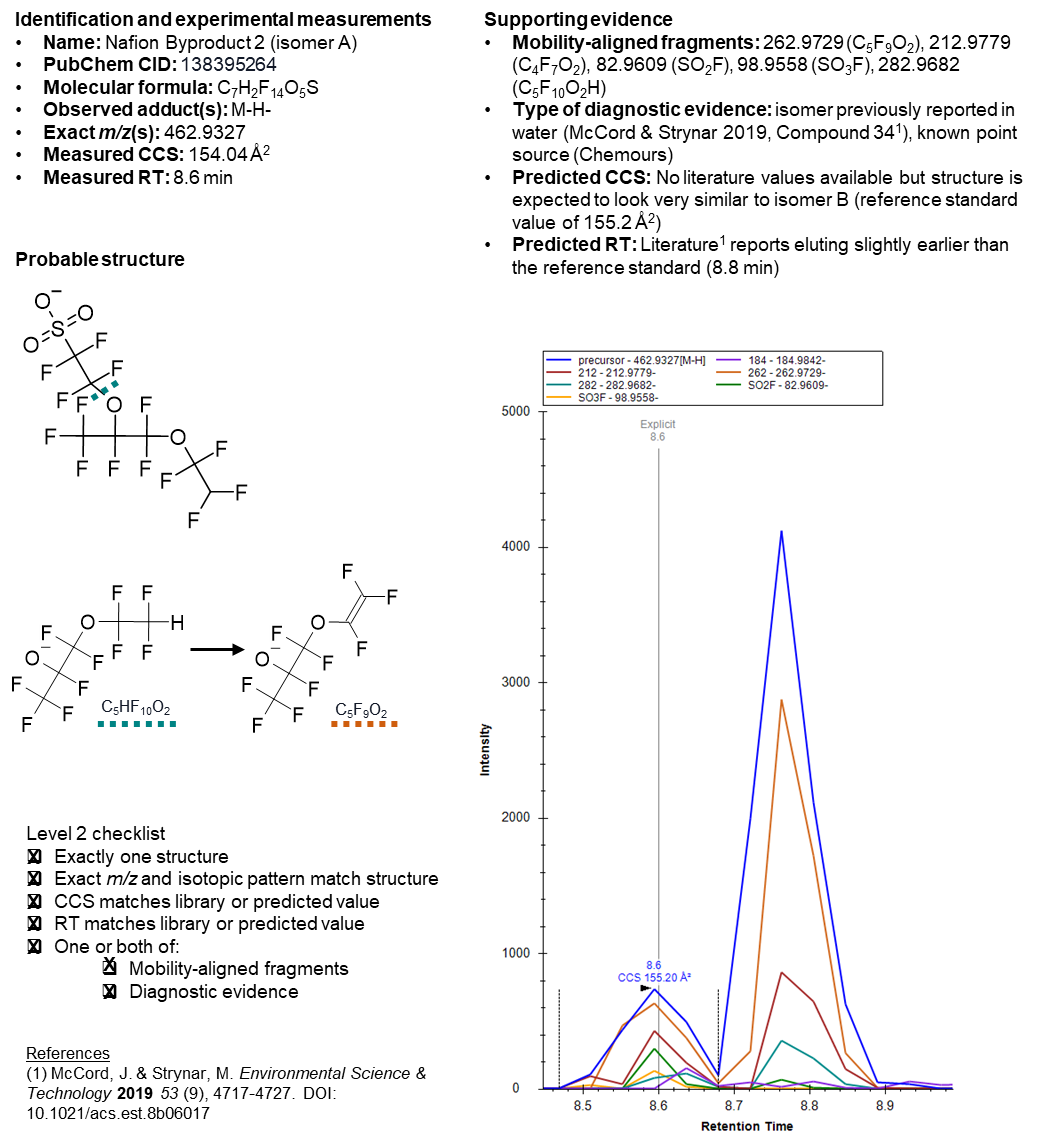

Fig. S10.

Supporting evidence for NB2 isomer A.

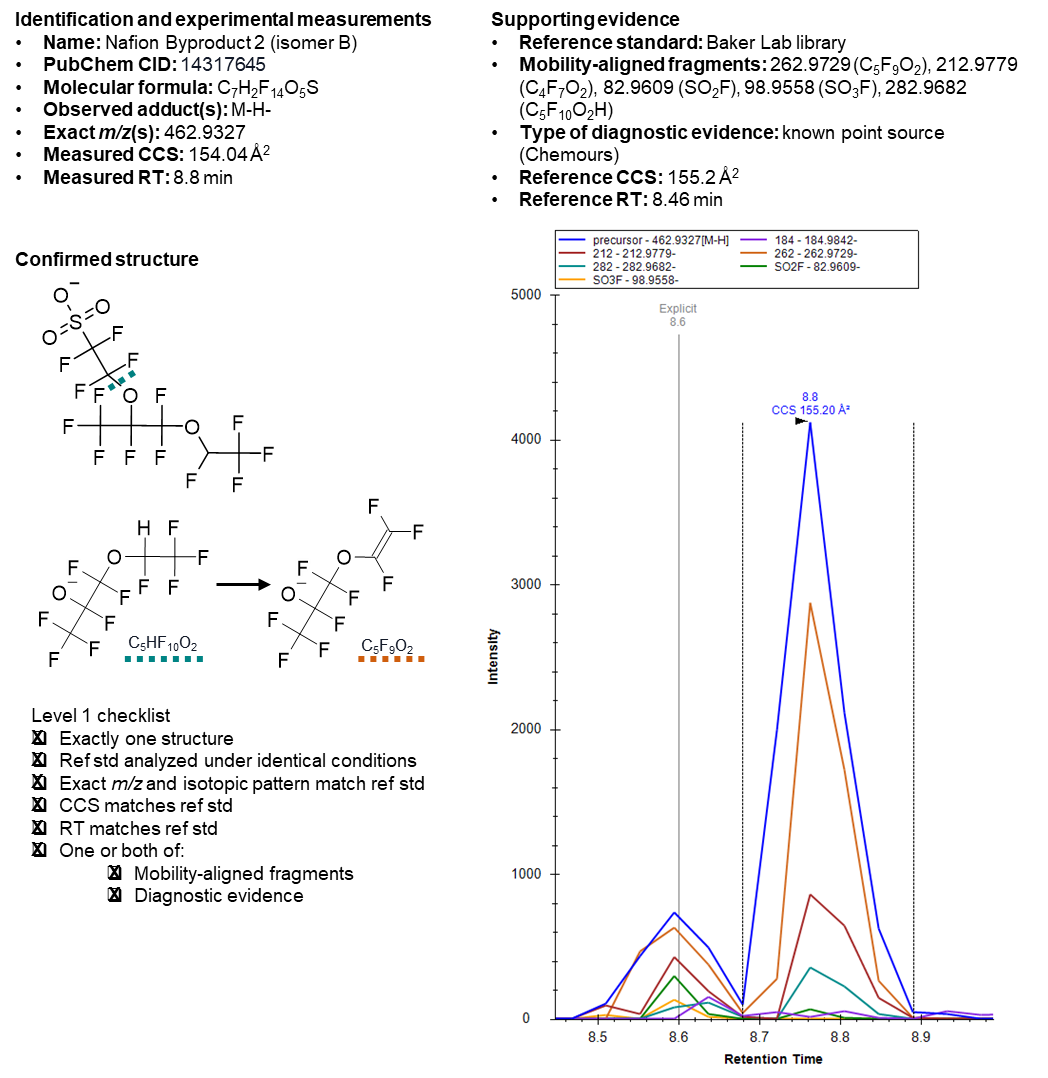

Fig. S11.

Supporting evidence for NB2 isomer B.

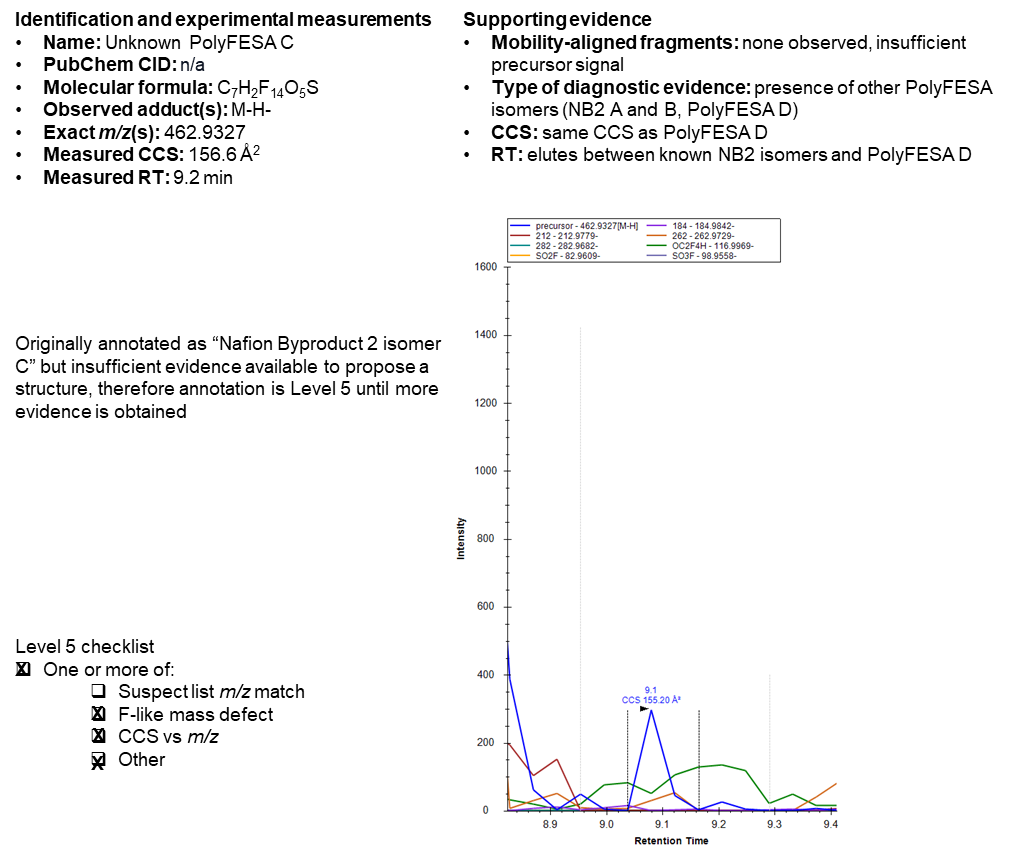

Fig. S12.

Supporting evidence for Unknown PolyFESA C.

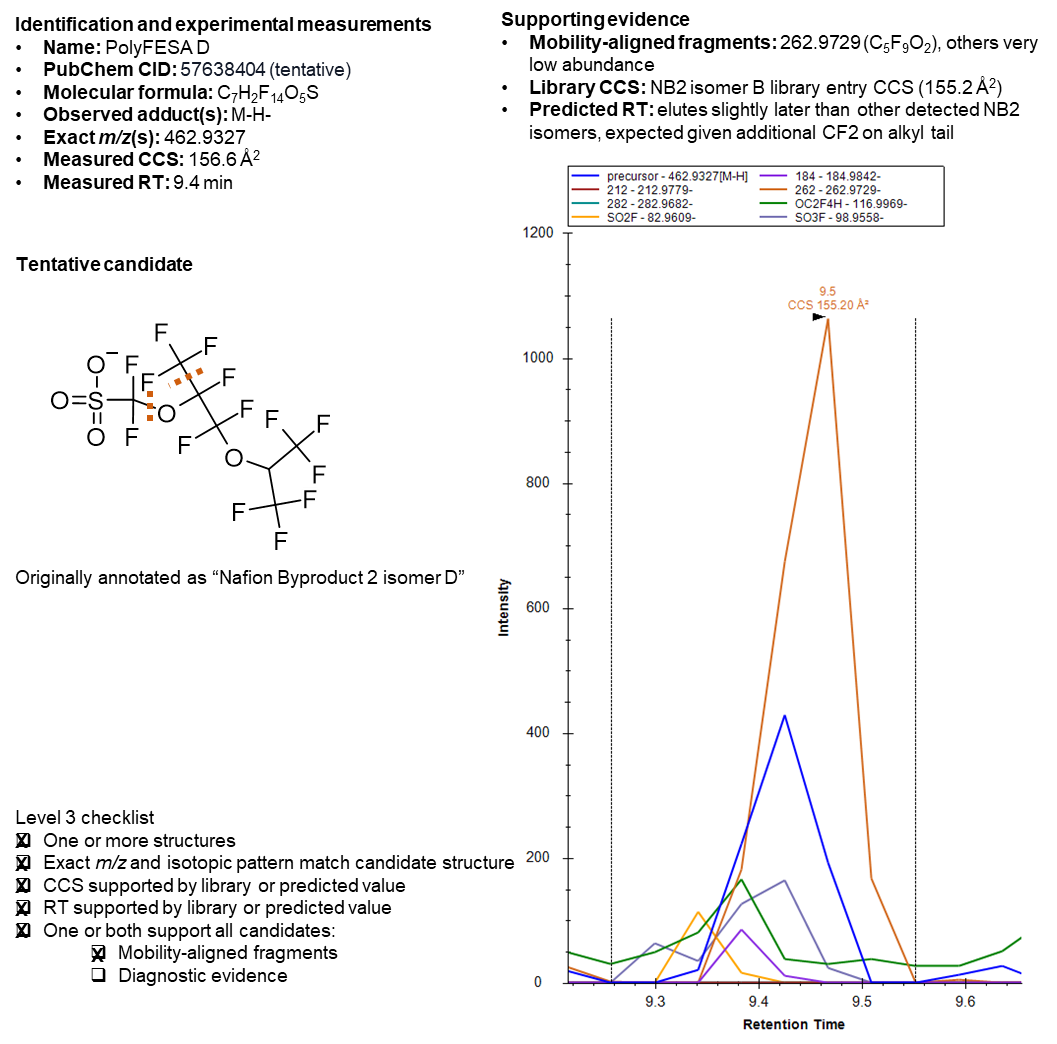

Fig. S13.

Supporting evidence for PolyFESA D.

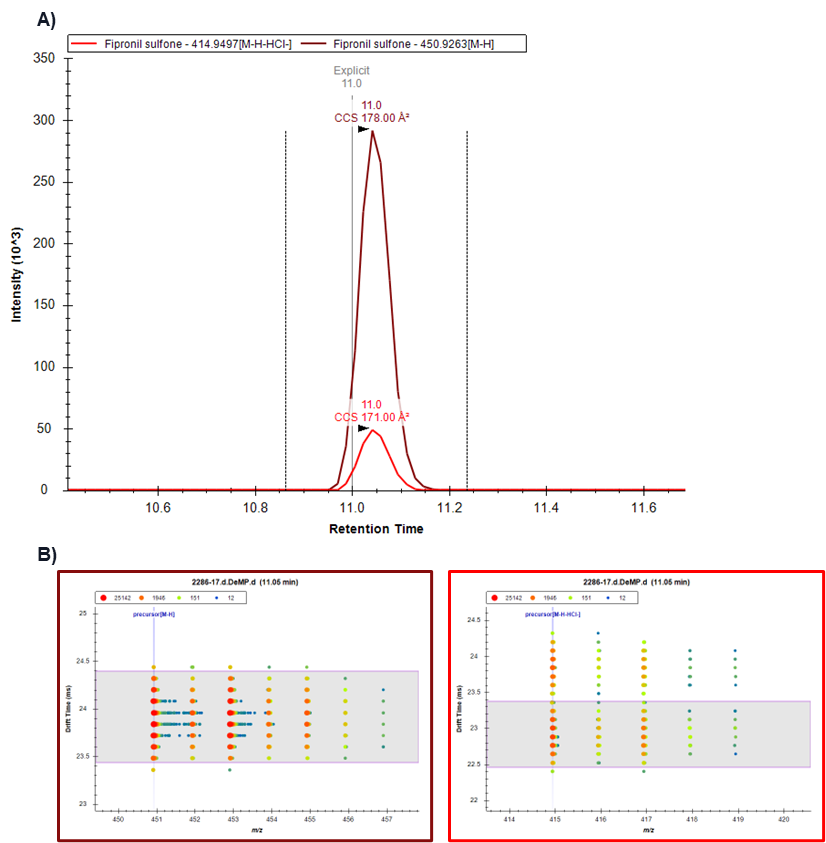

Fig. S14.

MS1-only data supporting fipronil sulfone. A) Extracted ion chromatogram (EIC) from a representative sample showing peaks for both M-H- and M-H-HCl- isomers. B) Nested drift spectra for both ions.

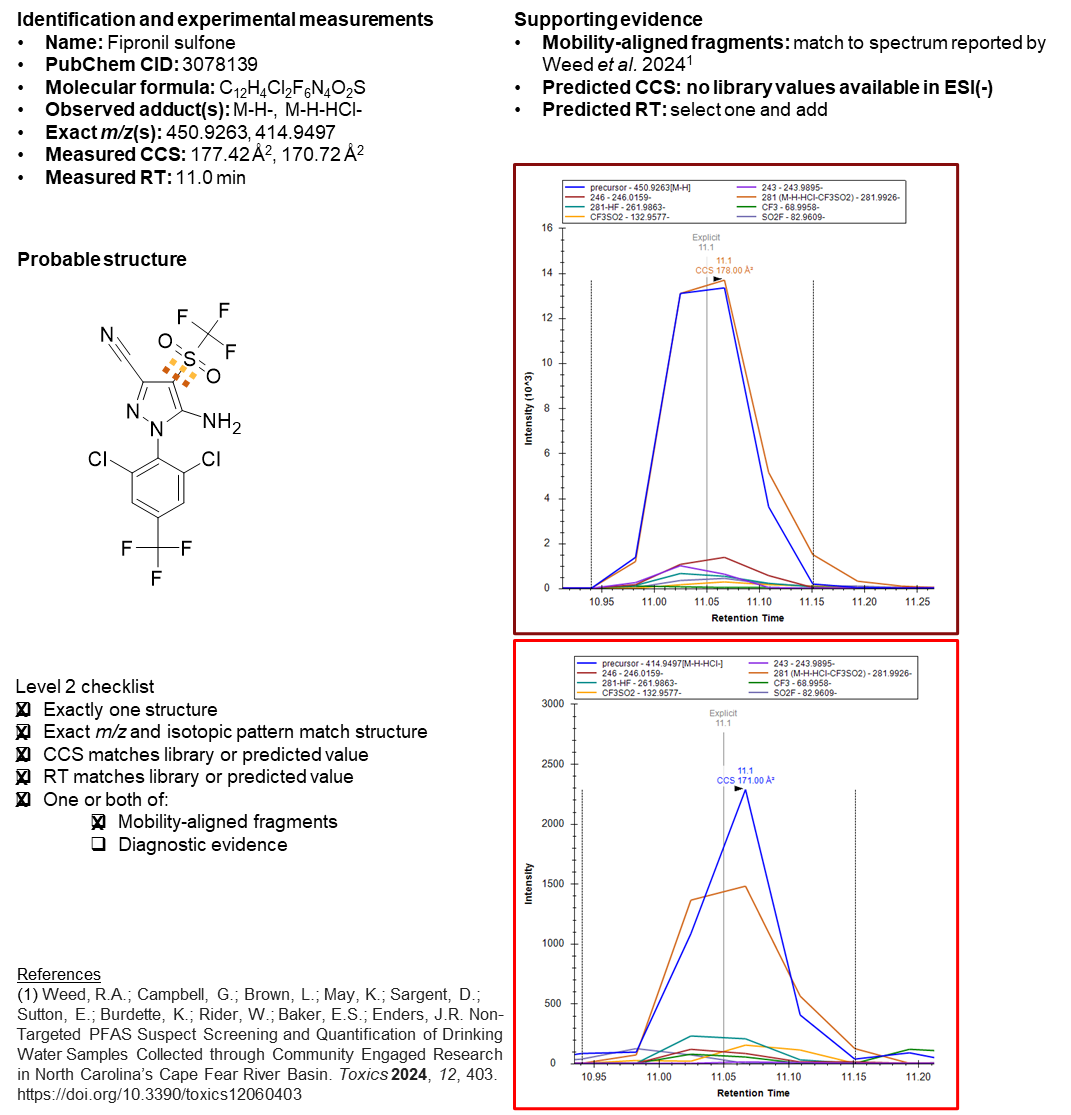

Fig. S15.

Supporting evidence for fipronil sulfone.

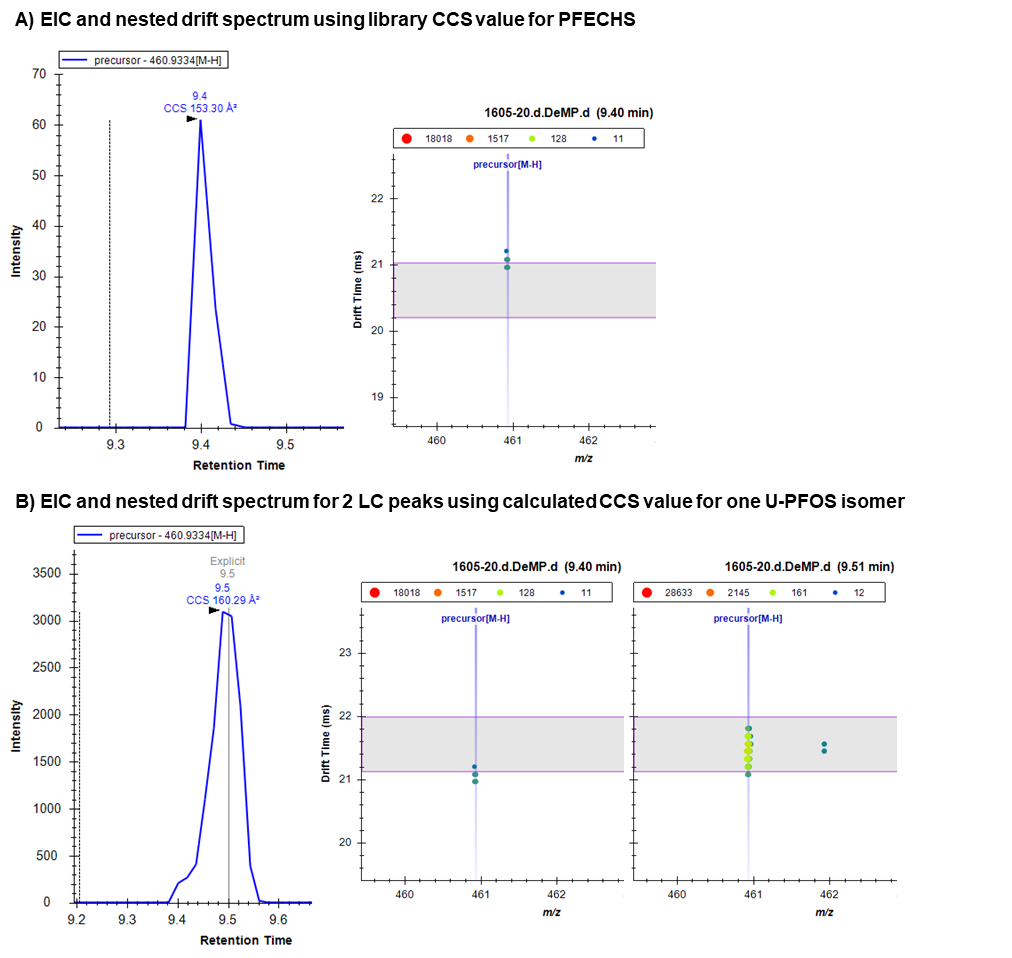

Fig. S16.

MS1-only data supporting U-PFOS. A) Extracted ion chromatogram (EIC) and nested drift spectrum using library CCS values for PFECHS, a structural isomer of U-PFOS. Horizontal bars show the drift window for PFECHS using a resolving power of 50. B) EIC using the calculated CCS for one U-PFOS isomer and drift spectra from two unresolved points along the LC peak showing presence of at least 2 isomers. The peak at 9.40 min is integrated in both A and B but clearly has a different CCS than either PFECHS or the isomer at 9.51 min.

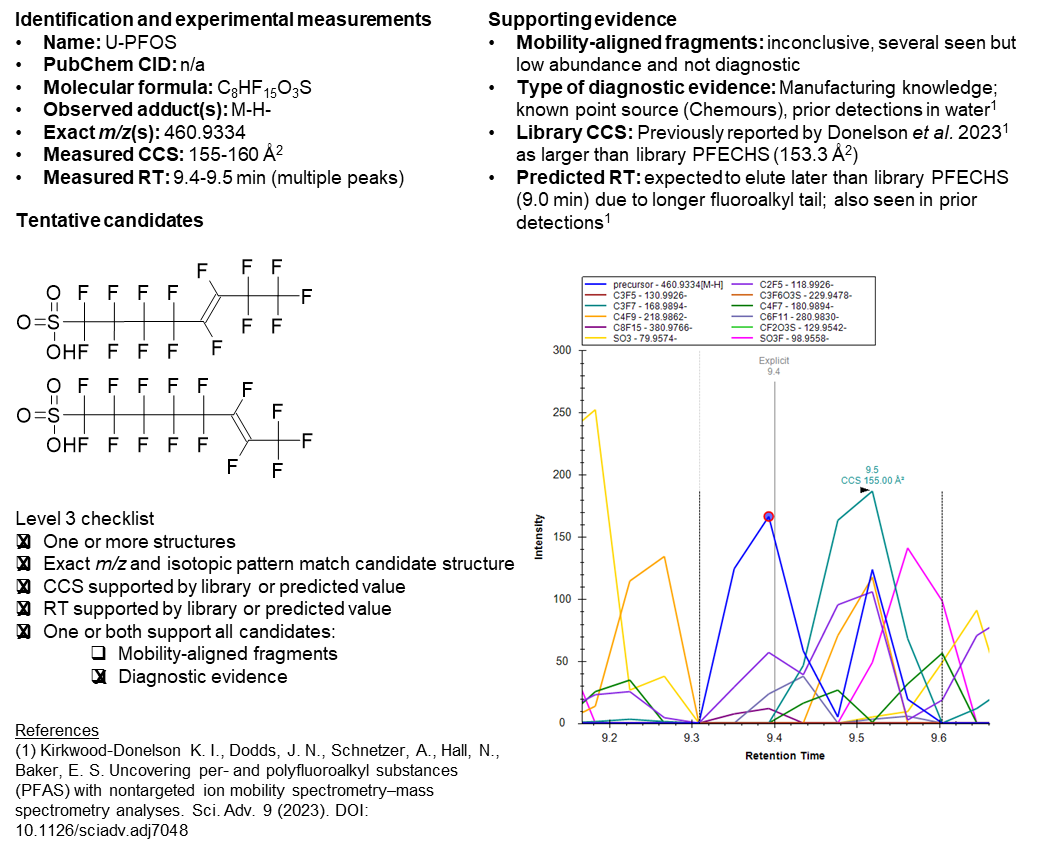

Fig. S17.

Supporting evidence for U-PFOS.

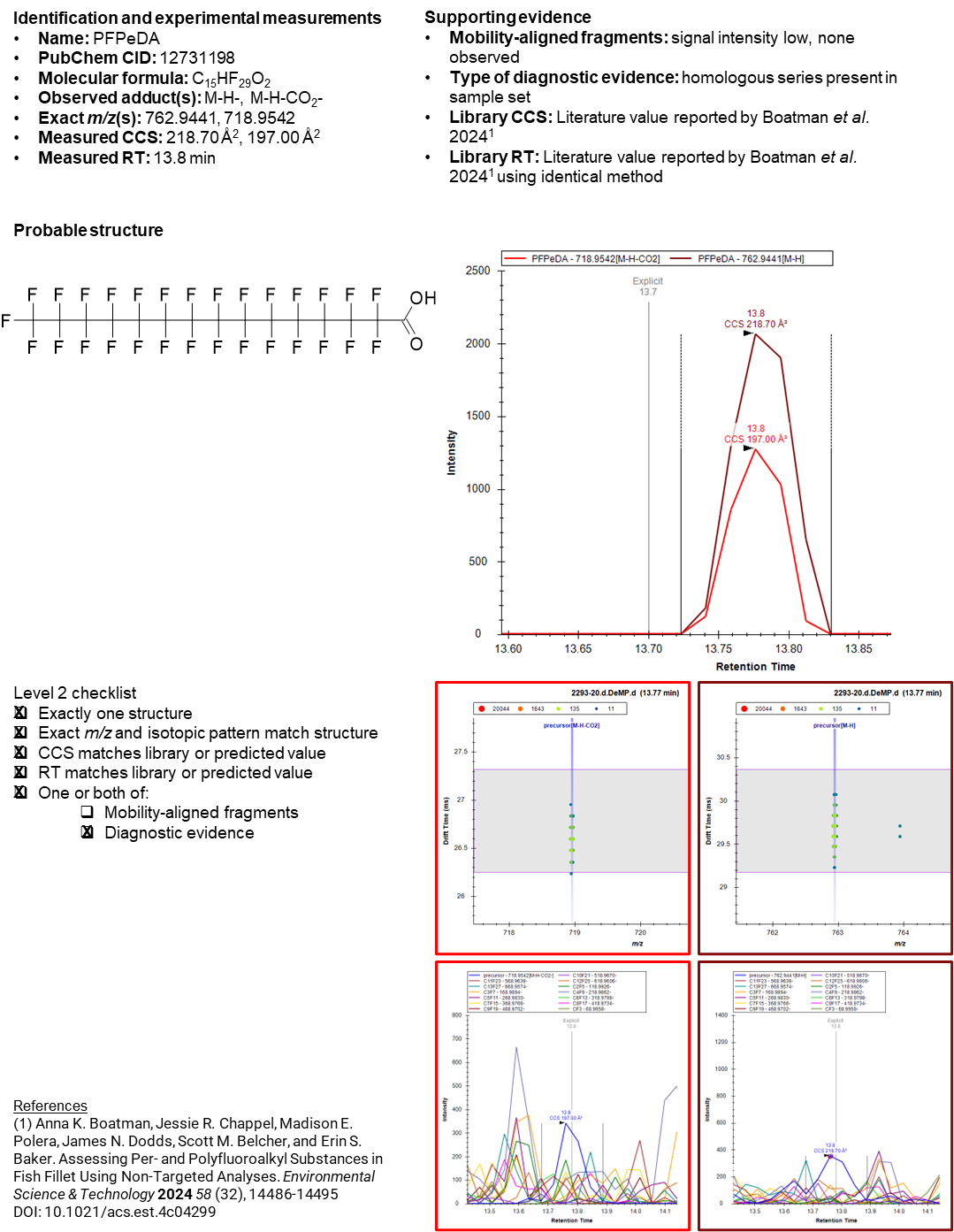

Fig. S18.

Supporting evidence for PFPeDA.

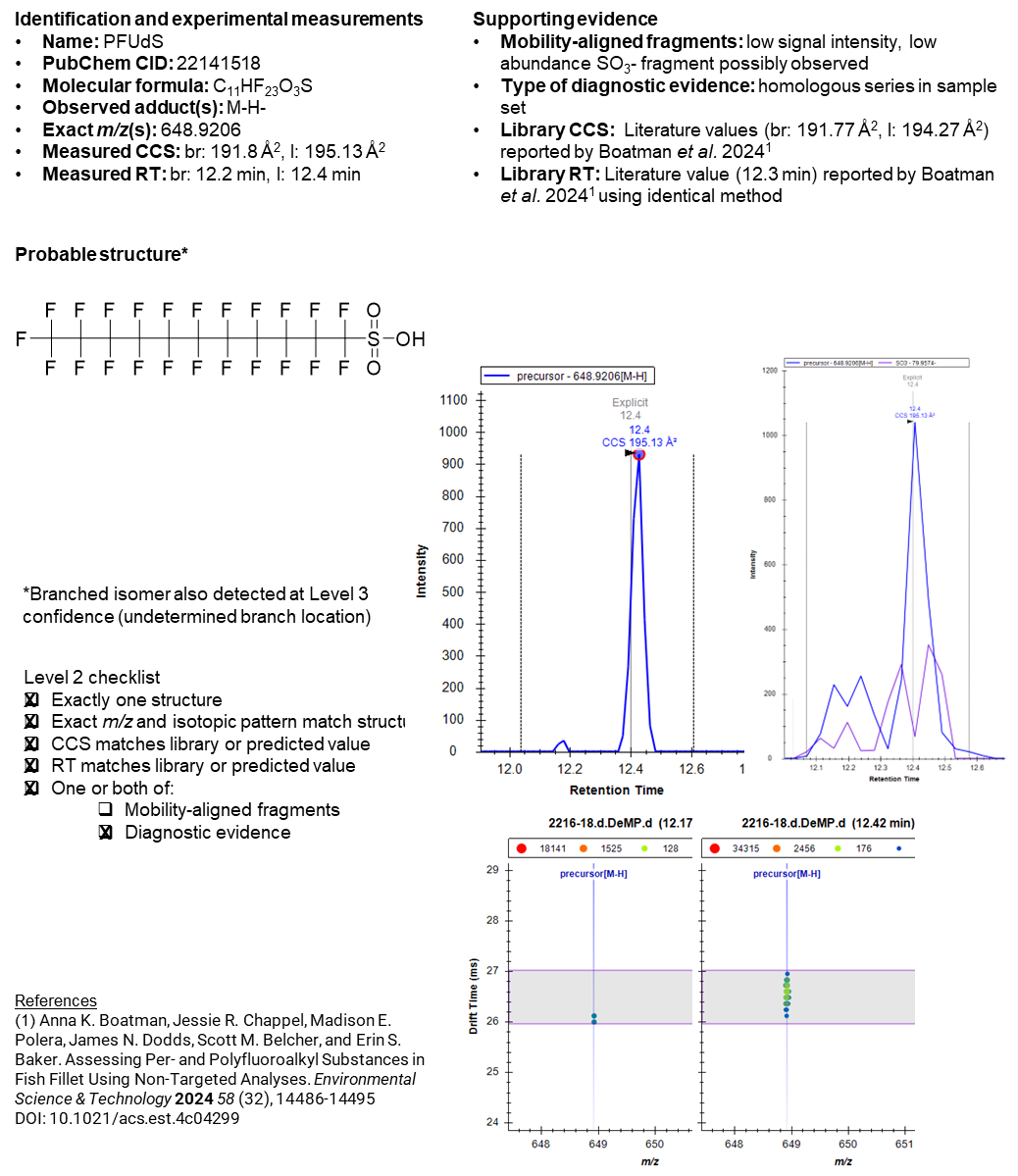

Fig. S19.

Supporting evidence for PFUdS.

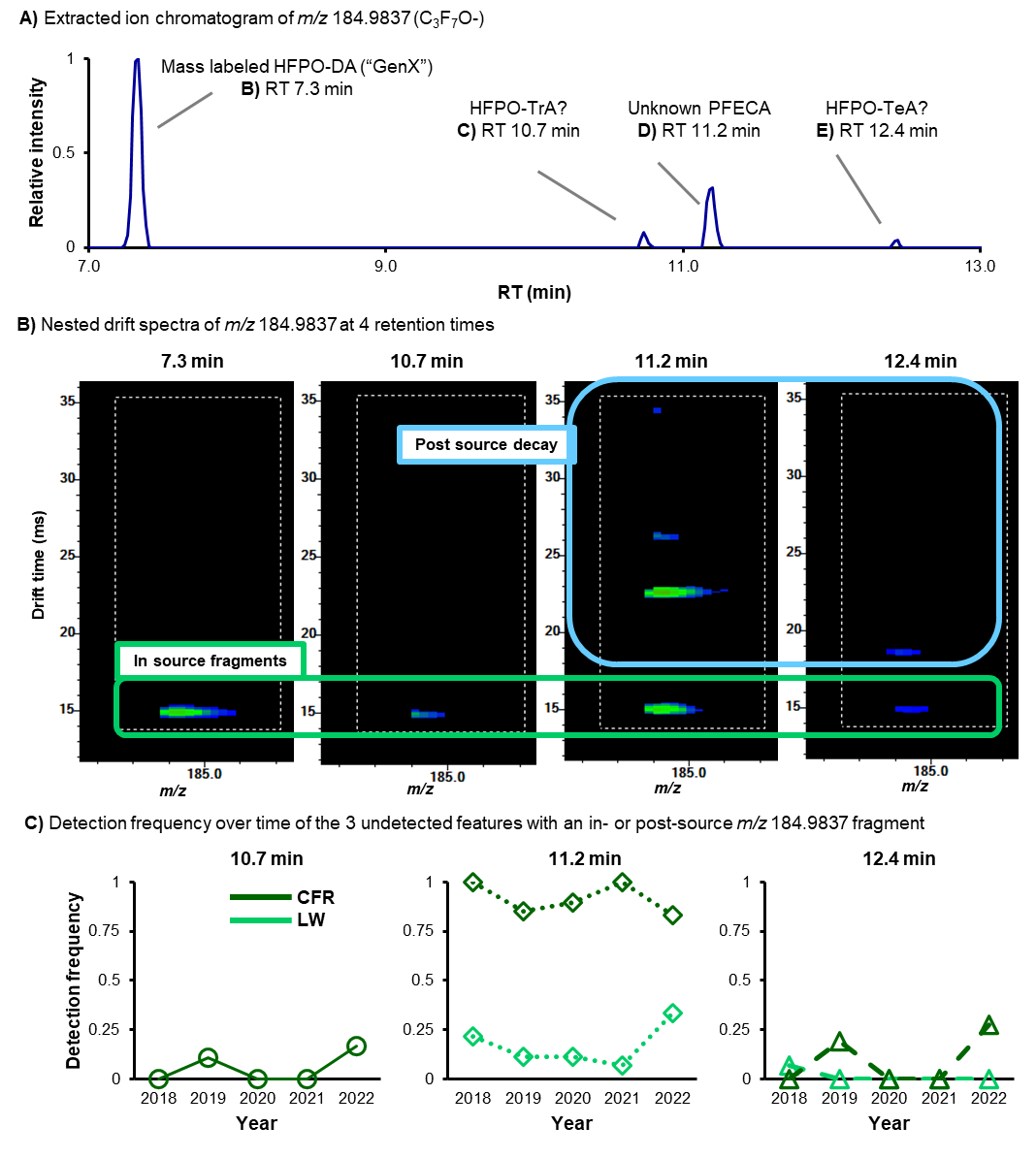

Fig. S20.

MS1-only fragmentation evidence of undetected PFECAs. A) EIC of C3F7O peaks B) Nested drift spectra. C) Detection frequency trends over time.

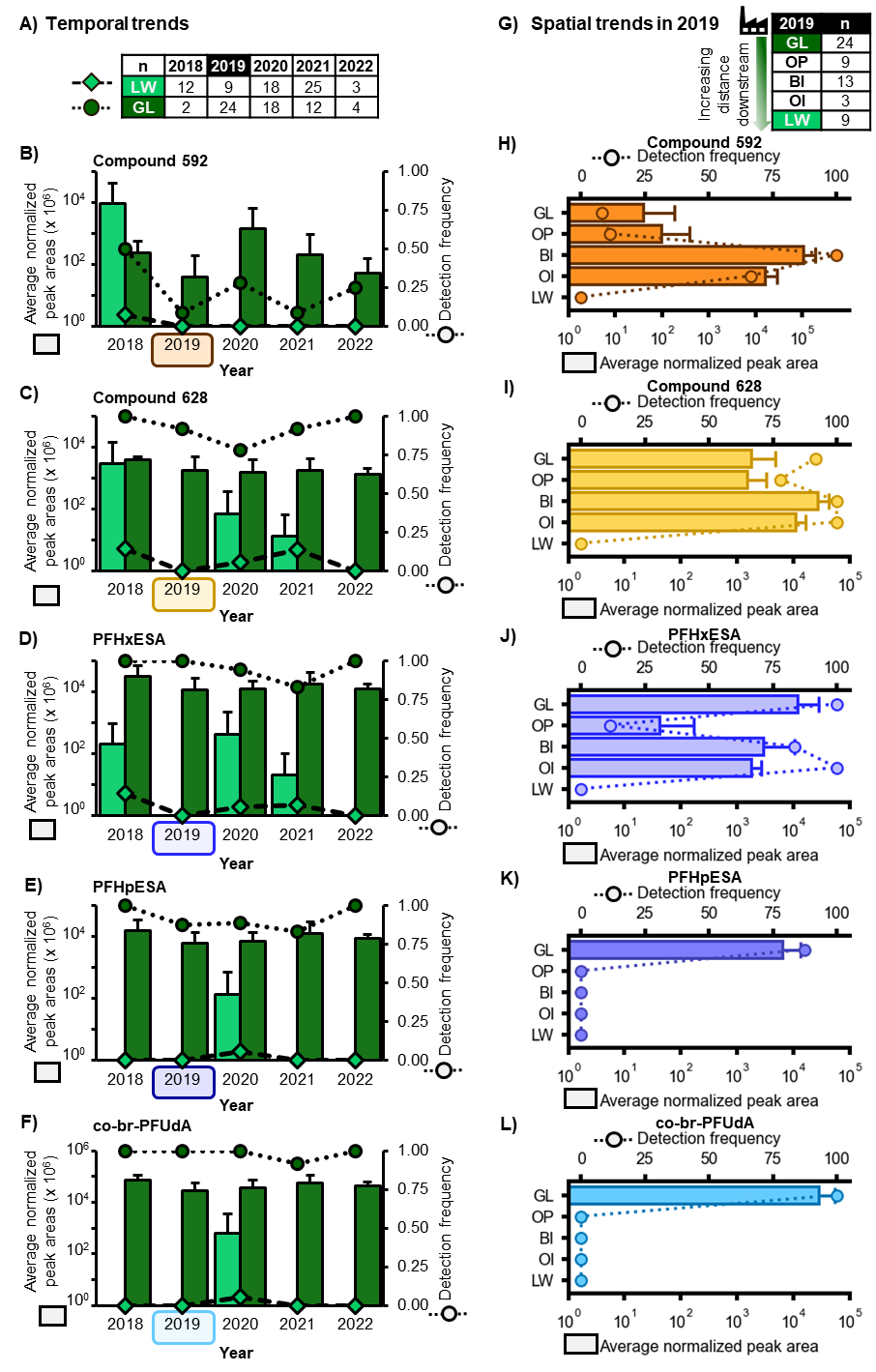

Fig. S21.

Preliminary temporal and spatial trend analysis for the 5 novel PFAS. GL: Greenfield Lake; LW: Lake Waccamaw; OP: Orton Pond; BI: Bald Head Island; OI: Oak Island. To minimize the effects of covariates, we selected the two sites with all 5 years of data available for the temporal trend analysis (Greenfield Lake in the CFR and the nearby reference site, LW). For the spatial trend analysis, we selected all sites with n ≥ 3 alligators from 2019 (the year with the most data available from different sites), including GL, LW, and 3 additional sites within the CFR: Orton Pond (OP), Oak Island (OI), and Bald Head Island (BI). None of the novel compounds were detected in FL alligators, which supports the North Carolina Chemours facility as a potential point source. A) Legend and number of alligators per year for the two sites with all 5 years of data available. B-F) Temporal trends in relative abundance (bar chart, left y-axis) and detection frequency (line graph with markers, right y-axis) over the 5 year study. G) Number of alligators per site for 2019, the year with the most different sites sampled with an n > 3 alligators per site.H-L) Site trends in relative abundance (bar chart, bottom axis) and detection frequency (line graph with markers, top axis). Sites are shown in order of distance downstream of Chemours.

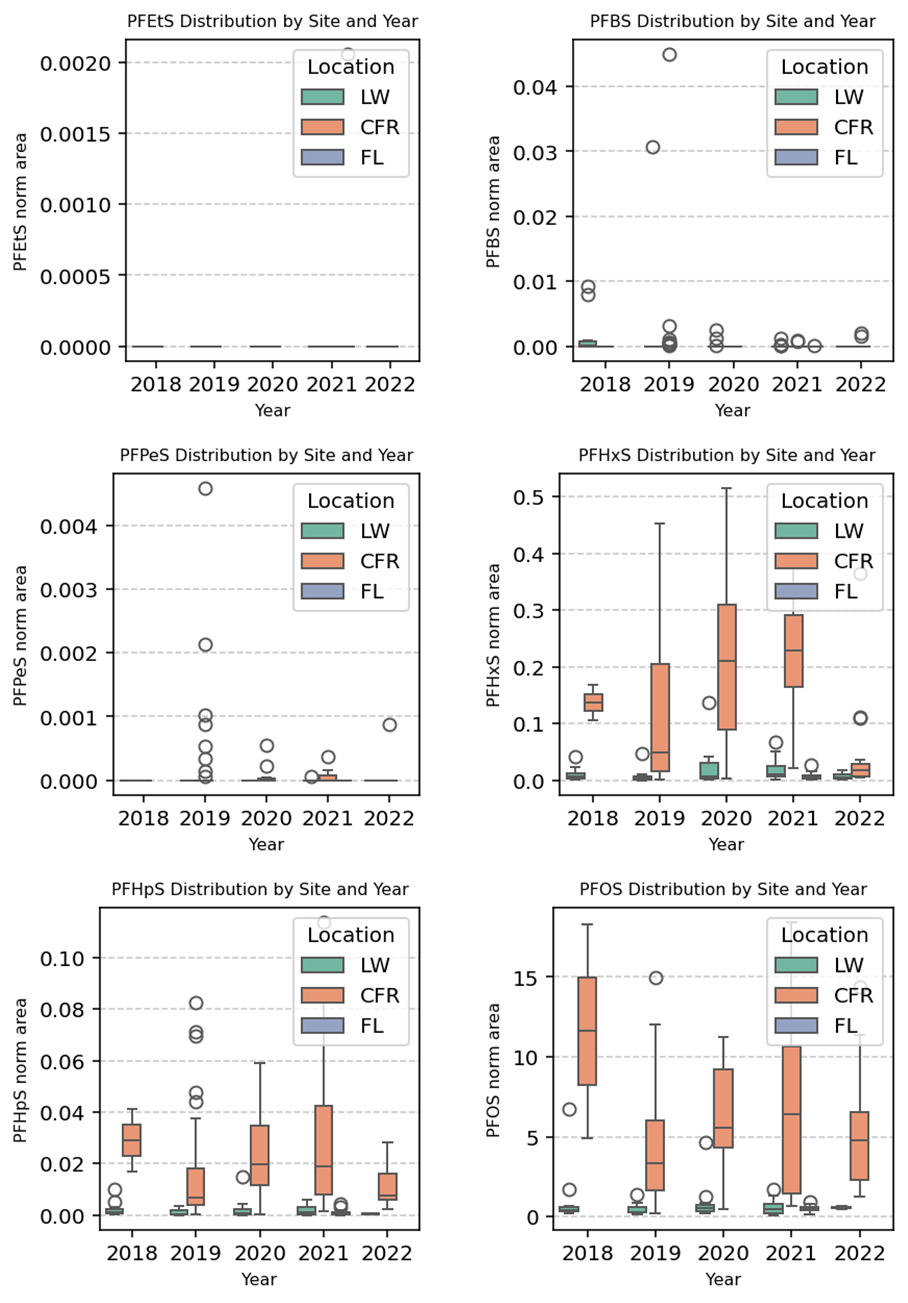

Fig. S22.

Relative abundance trends for PFSAs (PFEtS through PFOS). Box plots showing median and range of relative abundance for all detected PFSAs (PFEtS through PFOS) by watershed and year. Relative abundance is defined as the normalized peak area (total peak area divided by area of a surrogate standard).

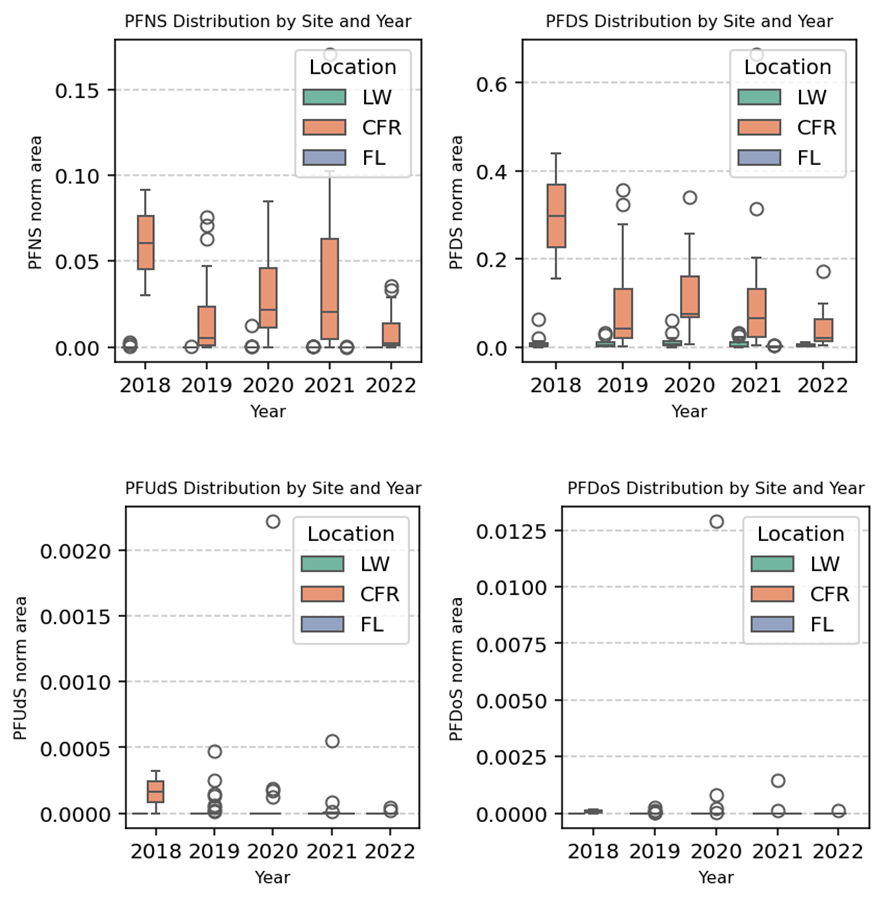

Fig. S23.

Relative abundance trends for PFSAs (PFNS through PFDoS). Box plots showing median and range of relative abundance for all detected PFSAs (PFNS through PFDoS) by watershed and year. Relative abundance is defined as the normalized peak area (total peak area divided by area of a surrogate standard).

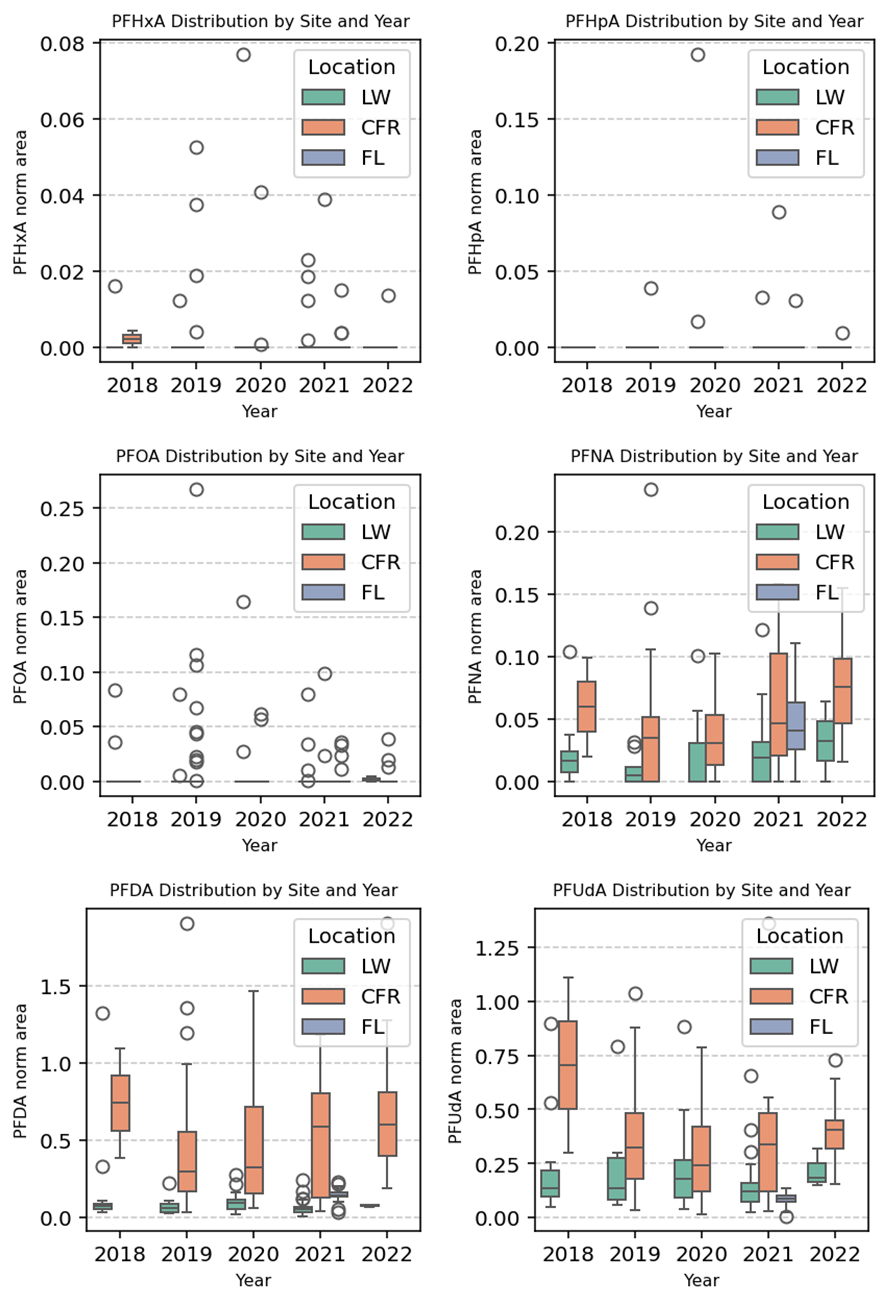

Fig. S24.

Relative abundance trends for PFCAs (PFHxA through PFUdA). Box plots showing median and range of relative abundance for all detected PFCAs (PFHxA through PFUdA) by watershed and year. Relative abundance is defined as the normalized peak area (total peak area divided by area of a surrogate standard).

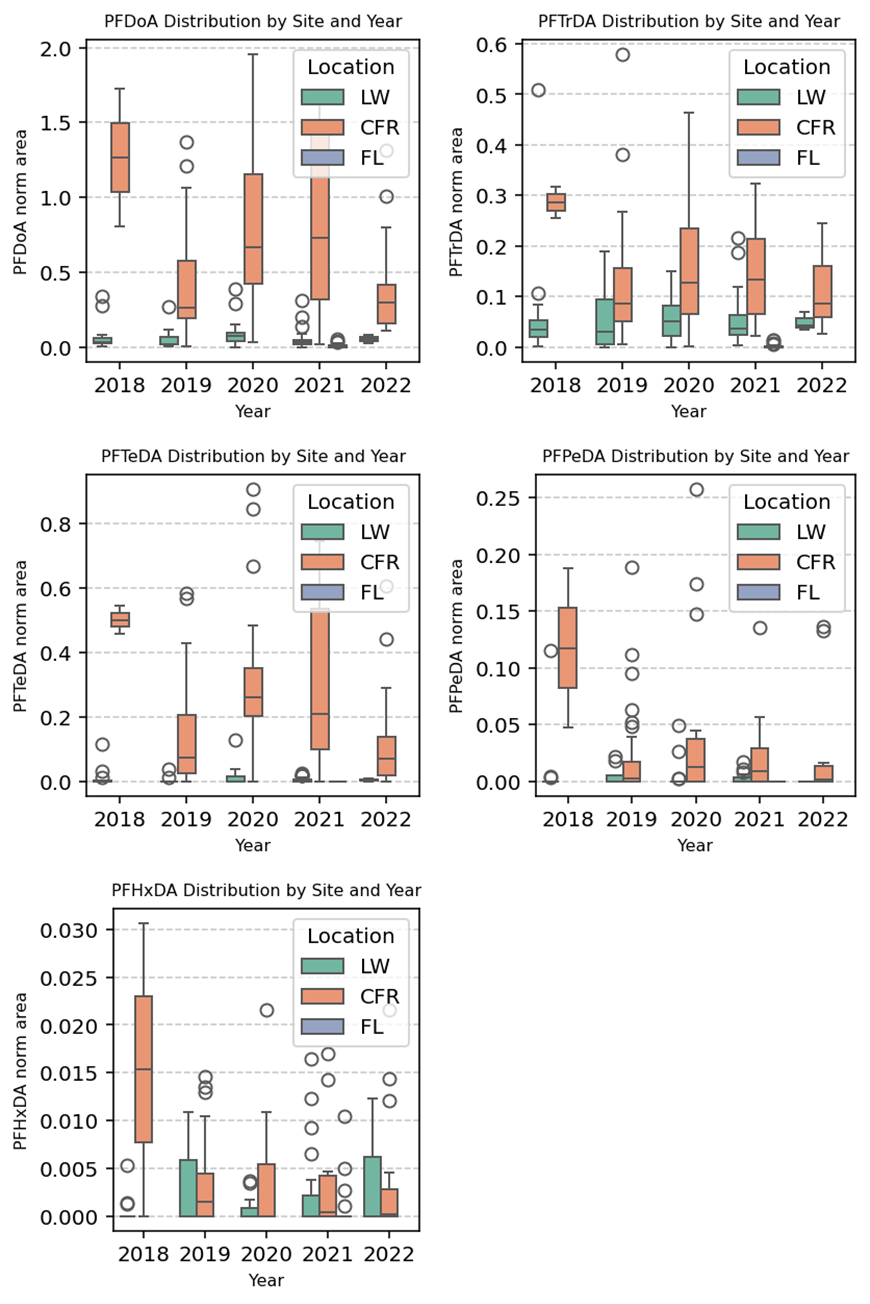

Fig. S25.

Relative abundance trends for PFCAs (PFDoA through PFHxDA). Box plots showing median and range of relative abundance for all detected PFCAs (PFDoA through PFHxDA) by watershed and year. Relative abundance is defined as the normalized peak area (total peak area divided by area of a surrogate standard).

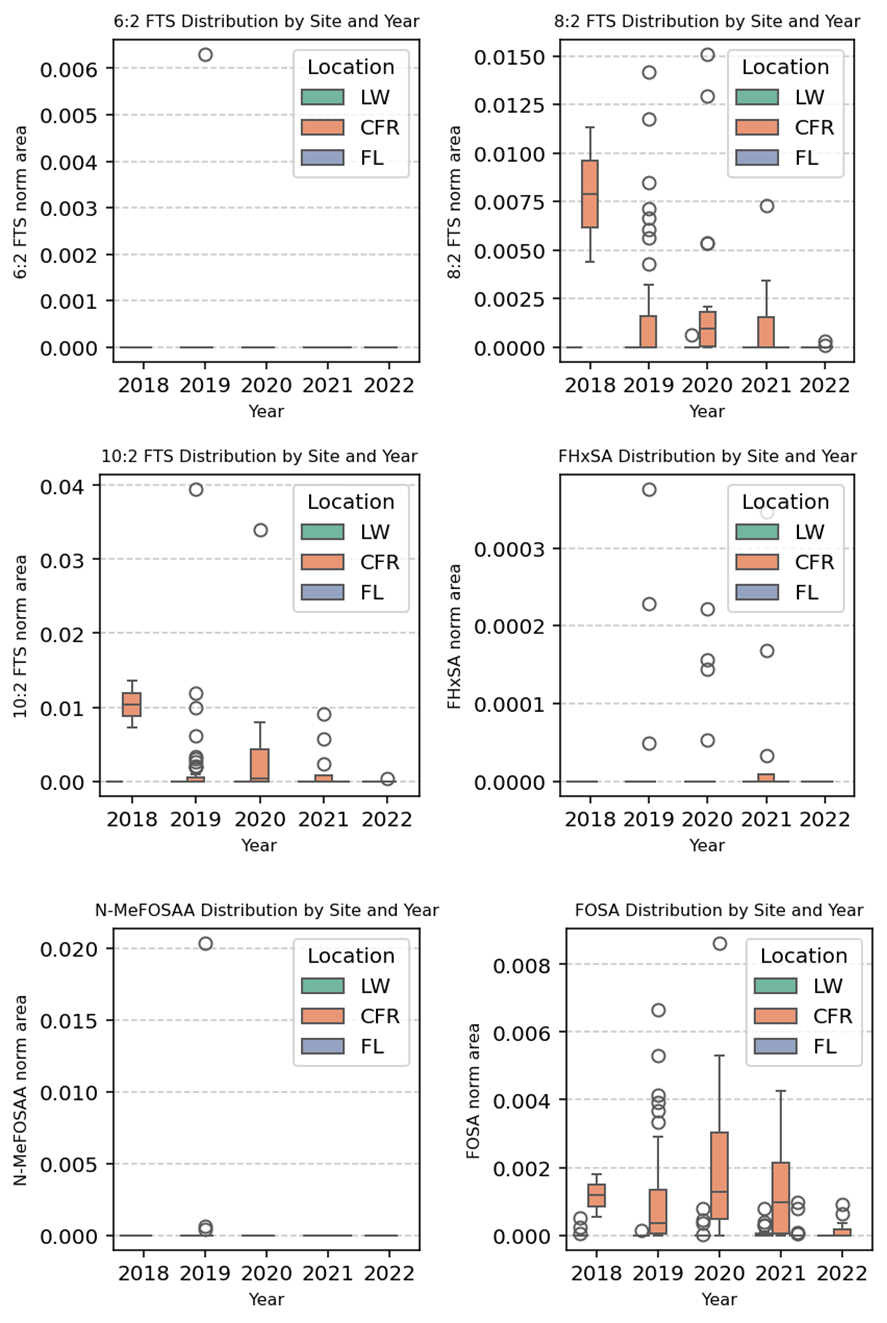

Fig. S26.

Relative abundance trends for FTSs and PFASAs. Box plots showing median and range of relative abundance for all detected fluorotelomer sulfonates (FTS) and perfluoroalkyl sulfonamides (FASAs) by watershed and year. Relative abundance is defined as the normalized peak area (total peak area divided by area of a surrogate standard).

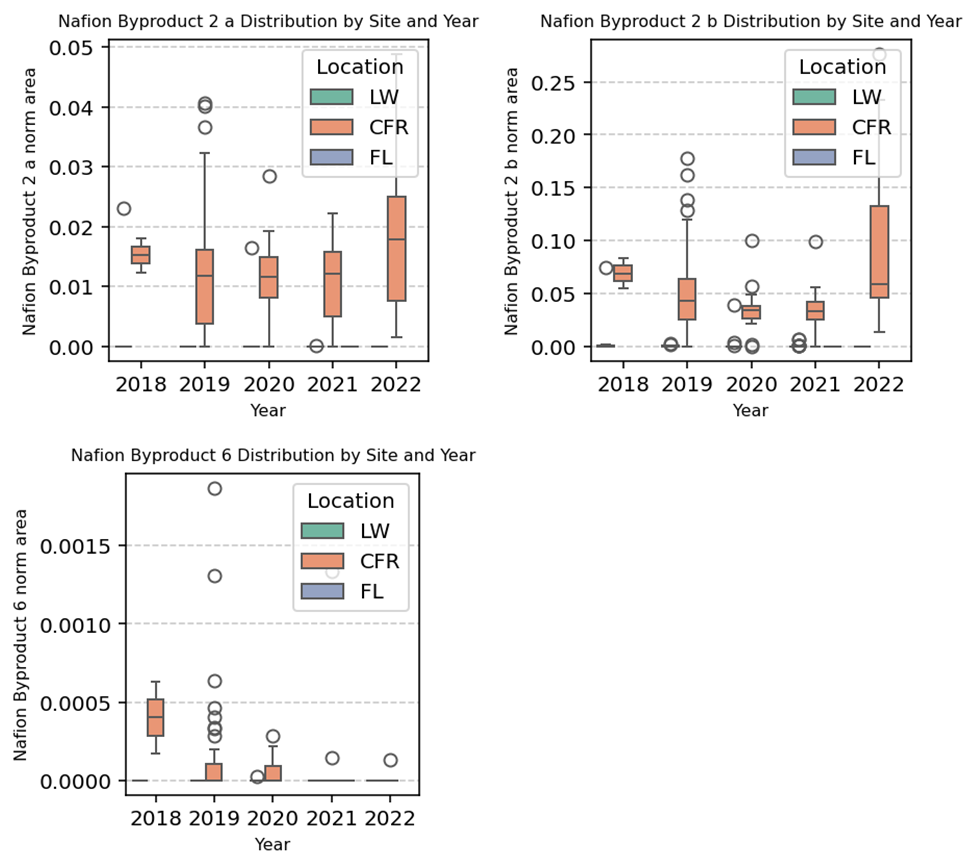

Fig. S27.

Relative abundance trends for Nafion byproducts. Box plots showing median and range of relative abundance for the two previously known Nafion 2 byproducts and Nafion Byproduct 6 by watershed and year. Relative abundance is defined as the normalized peak area (total peak area divided by area of a surrogate standard).

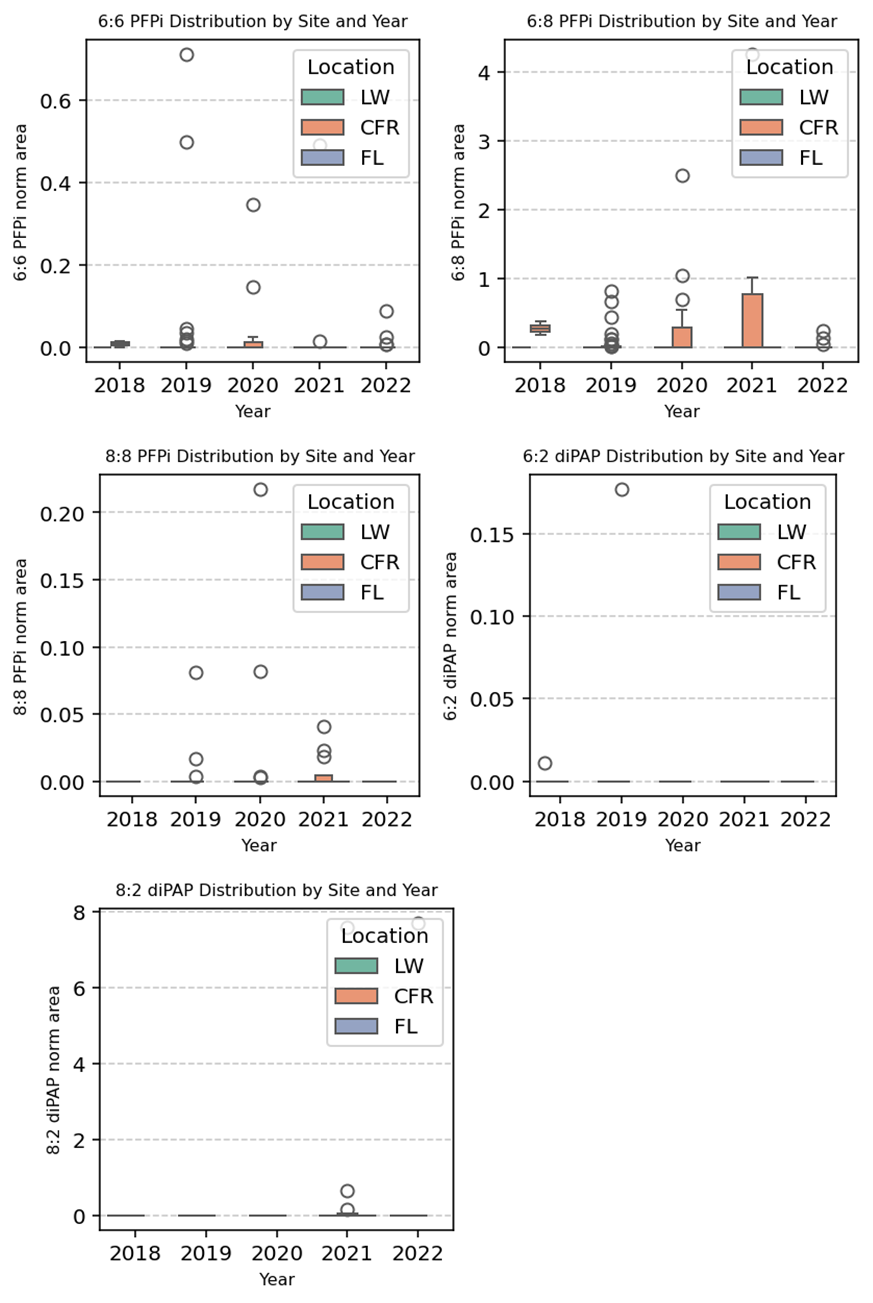

Fig. S28.

Relative abundance trends for PFPis and diPAPs. Box plots showing median and range of relative abundance for all detected perfluorophosphinic acids (PFPi) and polyfluoroalkyl phosphoric acid diesters (diPAPs) by watershed and year. Relative abundance is defined as the normalized peak area (total peak area divided by area of a surrogate standard).

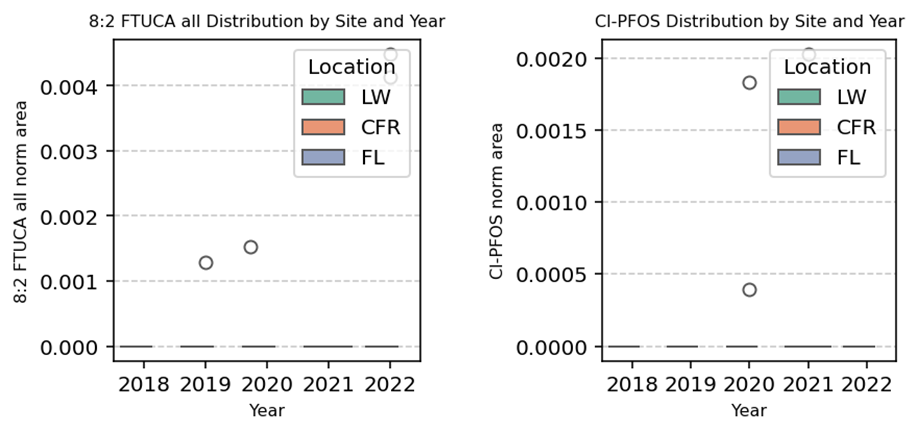

Fig. S29.

Relative abundance trends for 8:2 FTUCA and Cl-PFOS. Box plots showing median and range of relative abundance for 8:2 FTUCA and Cl-PFOS by watershed and year. Relative abundance is defined as the normalized peak area (total peak area divided by area of a surrogate standard).

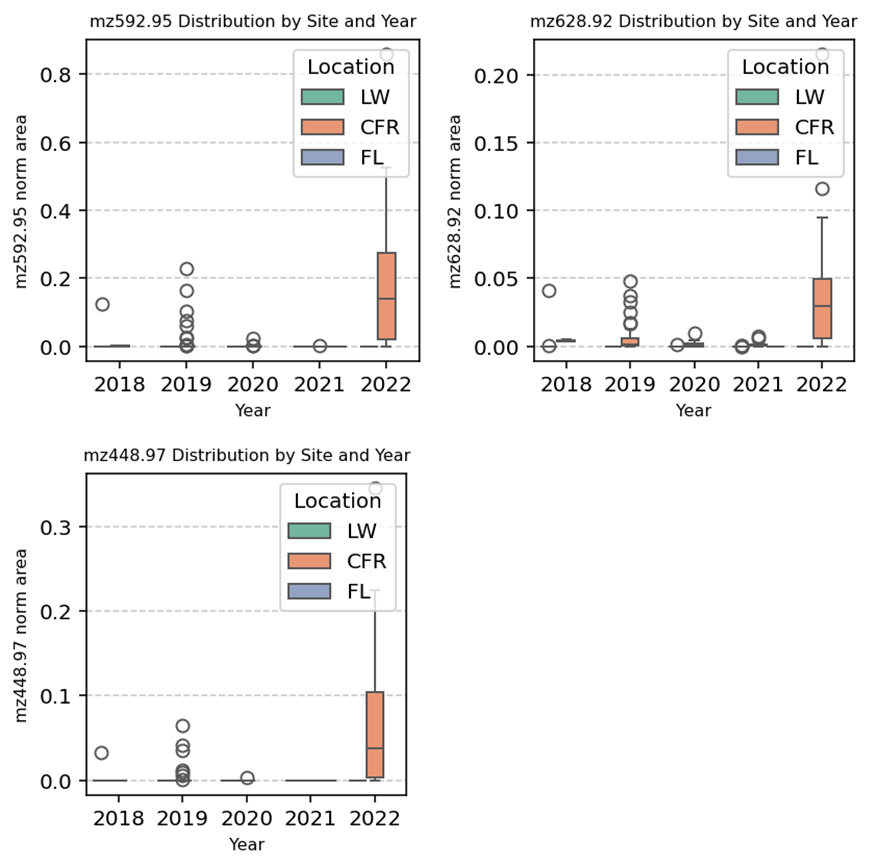

Fig. S30.

Relative abundance trends for novel triethers. Box plots showing median and range of relative abundance for the novel triether compounds by watershed and year. Relative abundance is defined as the normalized peak area (total peak area divided by area of a surrogate standard).

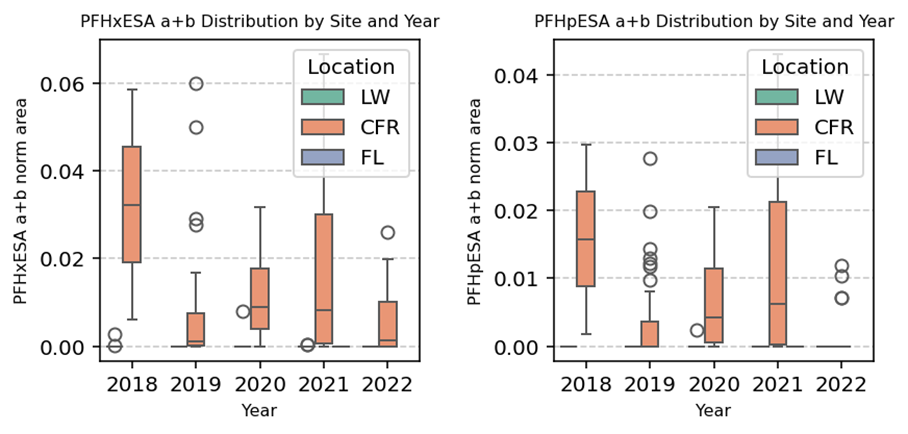

Fig. S31.

Relative abundance trends for novel PFAESAs. Box plots showing median and range of relative abundance for PFHxESA and PFHpESA by watershed and year. Peak areas of all detected isomers were summed. Relative abundance is defined as the normalized peak area (total peak area divided by area of a surrogate standard).

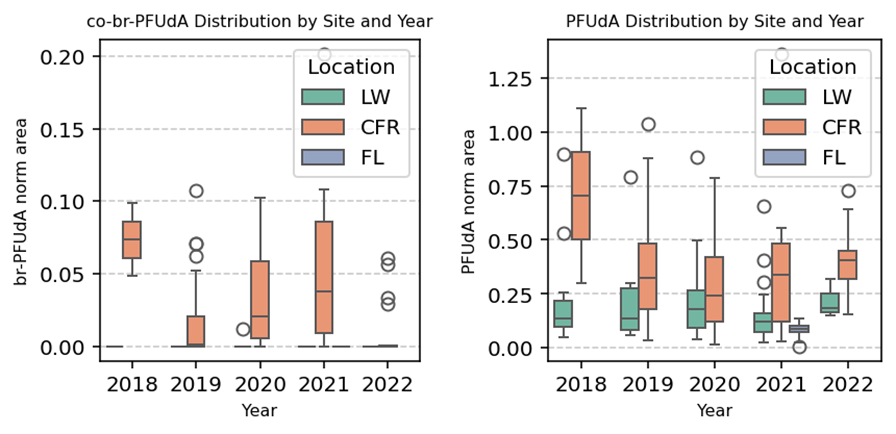

Fig. S32.

Relative abundance trends for PFUdA isomers. Box plots showing median and range of relative abundance for co-br-PFUdA and linear PFUdA by watershed and year. Relative abundance is defined as the normalized peak area (total peak area divided by area of a surrogate standard).

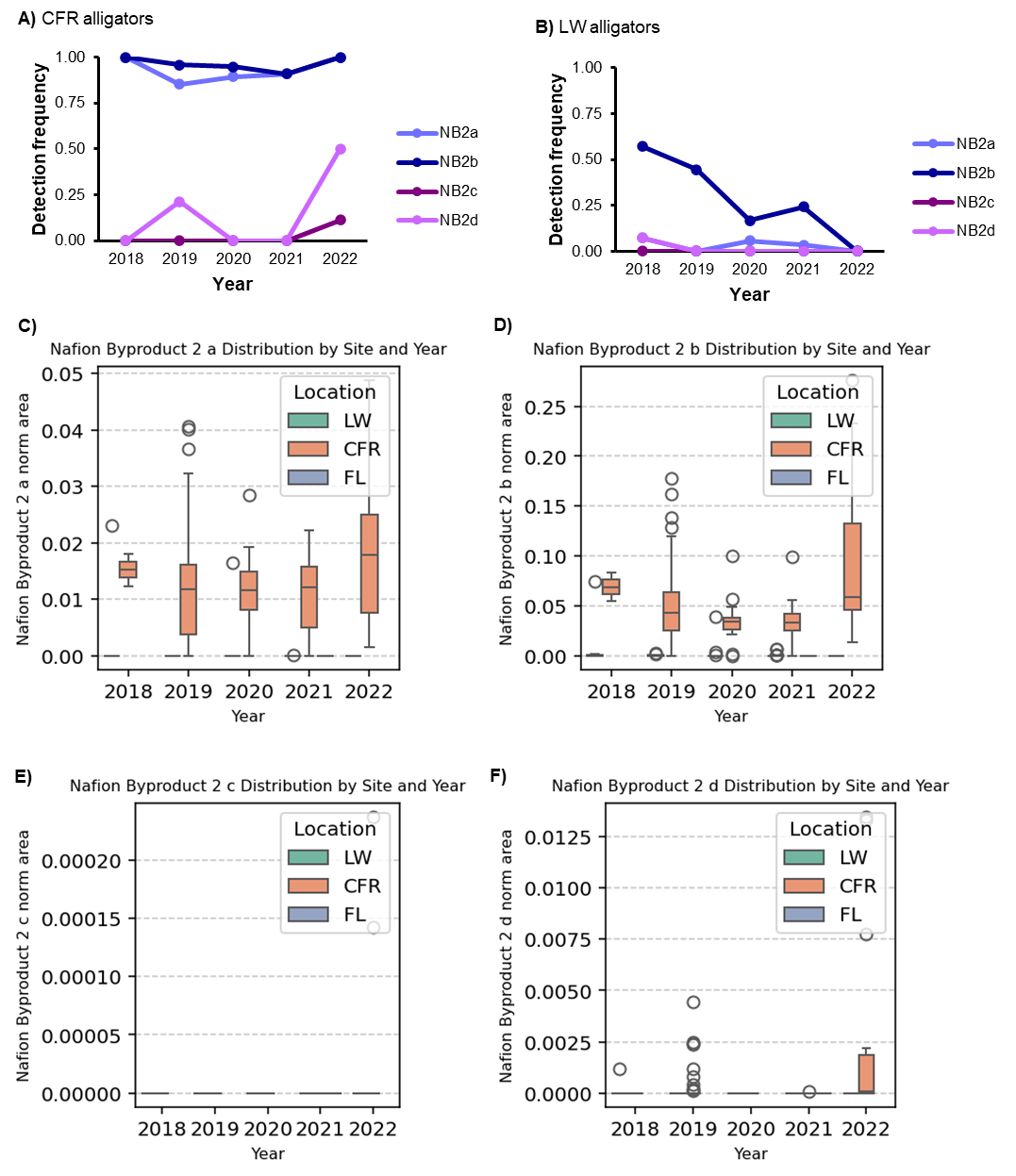

Fig. S33.

Relative abundance trends for PolyFESA isomers. Unknown PolyFESA C was originally annotated as “NB2 C” and PolyFESA D was originally annotated as “NB2 D”. A) Detection frequency in CFR alligators B) Detection frequency in LW alligators C-F) Box plots showing median and range of relative abundance for all detected NB2 isomers by watershed and year. Relative abundance is defined as the normalized peak area (total peak area divided by area of a surrogate standard).

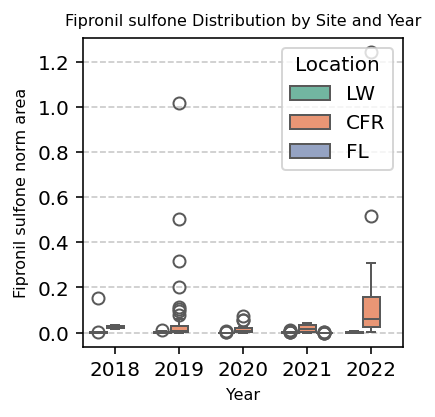

Fig. S34.

Relative abundance trends for fipronil sulfone. Box plots showing median and range of relative abundance for fipronil sulfone by watershed and year. Relative abundance is defined as the normalized peak area (total peak area divided by area of a surrogate standard).

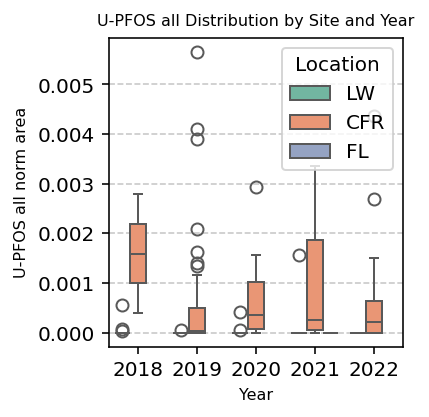

Fig. S35.

Relative abundance trends for U-PFOS. Box plots showing median and range of relative abundance for U-PFOS by watershed and year. Relative abundance is defined as the normalized peak area (total peak area divided by area of a surrogate standard).

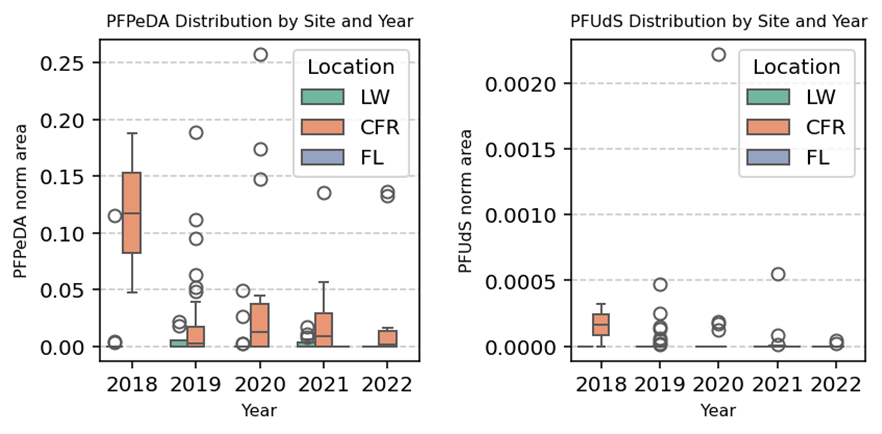

Fig. S36.

Relative abundance trends for PFPeDA and PFUdS. Box plots showing median and range of relative abundance for PFPeDA and PFUdS by watershed and year. Relative abundance is defined as the normalized peak area (total peak area divided by area of a surrogate standard).

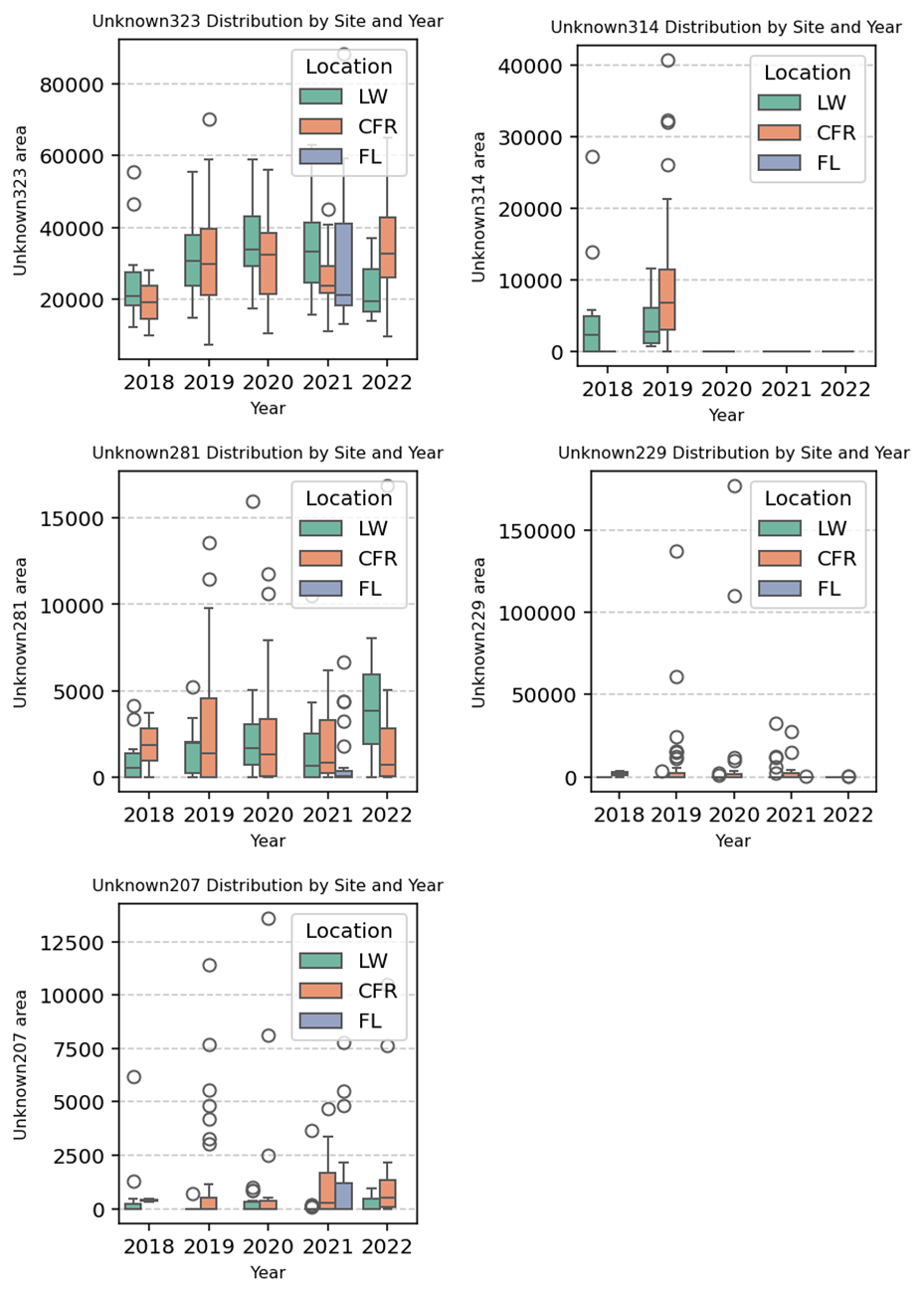

Fig. S37.

Peak area trends for the unidentified features of interest. Box plots showing median and range of peak areas for the unidentified features of interest by watershed and year.

Data S1. (separate file)

Alligator sampling locations and GPS coordinates.

Data S2. (separate file)

Data for individual alligator sample collection including pit tag number, GPS coordinates, collection and release times, sex, total length, snout vent length, and tail girth.

Data S3. (separate file)

Skyline transition list for all molecules evaluated in alligator samples.

Data S4. (separate file)

Skyline transition list for all detected molecules in alligator plasma samples, including surrogate standard used for internal standard peak area normalization.

Data S5. (separate file)

Relative abundances (normalized peak areas) for all PFAS detected in each alligator plasma sample.

Data S6. (separate file)

Absolute quantification summary and quality control results for the 26 PFAS included in the calibration curve.

Data S7. (separate file)

Absolute quantification results for individual alligator plasma samples.

Data S8. (separate file)

Absolute quantification results for quality control and blank samples.
